# A diverse population of pericoerulear neurons controls arousal and exploratory behaviors

**DOI:** 10.1101/2022.06.30.498327

**Authors:** Andrew T. Luskin, Li Li, Xiaonan Fu, Madison M. Martin, Kelsey Barcomb, Kasey S. Girven, Taylor Blackburn, Bailey A. Wells, Sarah T. Thai, Esther M. Li, Akshay N. Rana, Rhiana C. Simon, Li Sun, Lei Gao, Alexandria D. Murry, Sam A. Golden, Garret D. Stuber, Christopher P. Ford, Liangcai Gu, Michael R. Bruchas

## Abstract

As the primary source of norepinephrine (NE) in the brain, the locus coeruleus (LC) regulates arousal, avoidance and stress responses^1,2^. However, how local neuromodulatory inputs control LC function remains unresolved. Here we identify a population of transcriptionally, spatially and functionally diverse GABAergic neurons in the LC dendritic field that receive distant inputs and modulate modes of LC firing to control global arousal levels and arousal-related processing and behaviors. We define peri-LC anatomy using viral tracing and combine single-cell RNA sequencing with spatial transcriptomics to molecularly define both LC-NE and peri-LC cell types. We identify several neuronal cell types which underlie peri-LC functional diversity using a series of complementary neural circuit approaches in behaving mice. Our findings indicate that LC and peri-LC neurons are transcriptionally, functionally, and anatomically heterogenous neuronal populations which modulate arousal and avoidance states. Defining the molecular, cellular, and functional diversity of the LC and peri-LC provides a road map for understanding the neurobiological basis of arousal, motivation and neuropsychiatric disorders.

## Main

The locus coeruleus (LC) is the primary source of norepinephrine (NE) across the central nervous system, and is widely recognized as a critical modulator of global arousal states, including global autonomic activity and wakefulness, as well as more granular cognitive processes, such as memory, decision-making and attention^1,2^. The LC has been implicated in having heterogenous roles in pain, avoidance and reinforcement learning, exploration, and feeding, which are likely modulated by distinct LC subpopulations with diverse morphology, projection patterns, and electrophysiological properties^3–10^. However, it remains unclear how LC ensembles and their activity dynamics are controlled locally. Long-range neuromodulatory inputs to the region potently modulate the LC, but many do not target LC neurons directly^11–15^. LC dendrites extend medially to form a pericoerulear zone (the peri-LC)^16^ that is believed to receive inputs from the medial prefrontal cortex (mPFC), dorsomedial and paraventricular hypothalamus, central amygdala, medial preoptic area, and dorsal raphe^11–13^. An uncharacterized population of GABAergic neurons in the peri-LC has been suggested to modulate physiological arousal by inhibiting the LC^11,17,18^. We hypothesized that the peri-LC^GABA^ population plays critical and important roles in coordinating LC activity and can influence arousal-related and avoidance behaviors by modulating LC^NE^ firing. Here, we report that peri-LC^GABA^ neurons directly inhibit the LC and reveal that these neurons increase their activity in behaving mice in response to salient stimuli, during exploration, and after feeding. Using a combination of optogenetic and chemogenetic manipulations, we found that stimulating peri-LC^GABA^ neurons potently decreases physiological arousal and exploration, while inhibiting them increases avoidance or anxiety-like behaviors. Complementary single-nucleus and spatial RNA transcriptomics uncovered the diverse features of both peri-LC and LC neuronal cell types, and these results informed further experiments which revealed differential neural responses and a sufficient role for neuropeptide-containing peri-LC^GABA^ subpopulations in altering arousal-related and avoidance behaviors. Together, our results reveal specialized roles for this group of locally projecting, inhibitory peri-LC neurons in tuning LC^NE^ activity to regulate arousal and avoidance states.

### Peri-LC^GABA^ neurons receive distant inputs and directly control LC activity

We first examined the anatomical properties of peri-LC^GABA^ neurons in the pericoerulear zone, the area medial to the LC and spanning the length of the LC dendritic field, from approximately 5.3 to 5.8 mm caudal to bregma^16^. Using Cre-inducible viral expression of enhanced yellow fluorescent protein tagged channelrhodopsin-2 (AAV5-DIO-ChR2-eYFP) and immunostaining of tyrosine hydroxylase (TH) in cleared tissue of vesicular GABA transporter (vGAT)-Cre mice, we found that peri-LC^GABA^ neurons form a dense web on the ventromedial face of the LC, where they intertwine with LC^NE^ dendrites (**Fig. 1a, Supplemental Video 1**). These peri-LC^GABA^ neurons follow the LC^NE^ dendritic field, which is primarily medial in anterior LC and more ventral in posterior LC, and extends 300-400 μm from the LC, consistent with the length of LC^NE^ dendrites measured in previous studies^19^. Notably, these GABAergic projections are almost exclusively local; after examining the whole brains of mice, only a single long-range projection was found outside of the LC region, within the lateral interpeduncular nucleus. Additionally, using monosynaptic rabies tracing onto peri-LC^GABA^ starter cells, we found diverse inputs to peri- LC^GABA^ neurons from distant brain regions including the bed nucleus of the stria terminalis (BNST), the central amygdala (CeA), the lateral hypothalamus (LH), the medial cerebellar nucleus (Med), and numerous brainstem areas (**Extended Data Fig. 1a, Supplementary Table S1**), consistent with previous findings^18^, and increased our confidence in this area as a target of affective-motivational centers and ascending signals from arousing stimuli. We also identified inputs from pontine nuclei including the parabrachial nucleus, a hub for sensory information including pain, feeding control, and alarm signals^20^, and the pedunculotegmental and laterodorsal tegmental nuclei, implicated in attention, sleep-wake cycles, and reward^21^.

**Fig. 1.**
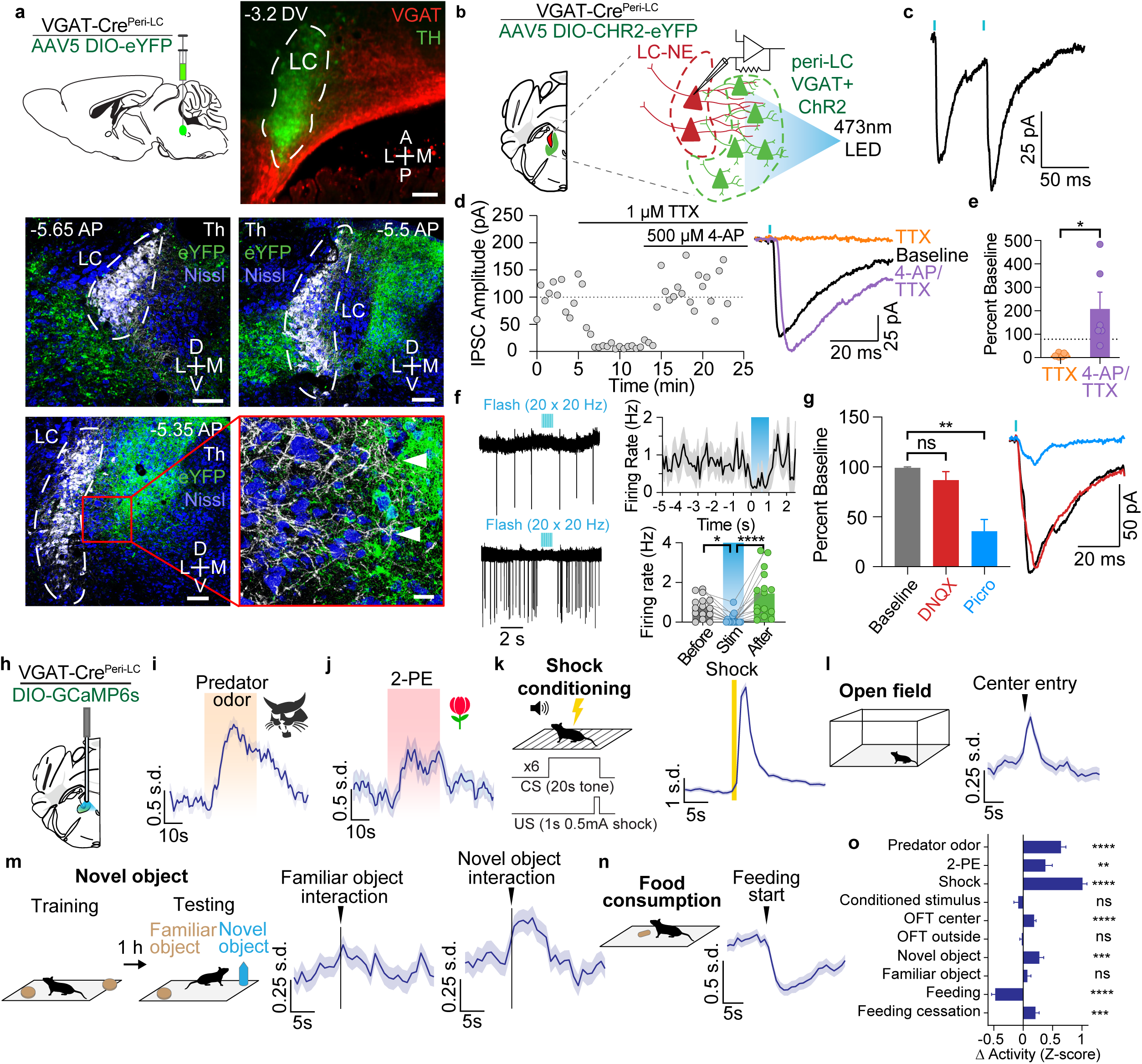
*In vitro* and *in vivo* identification of the peri-LC GABAergic neural network. **(a)** Schematic of viral injection in vGAT-Cre mice is shown on top left. Light-sheet image of cleared mouse brain immunostained for TH (green) and eYFP-expressing GABAergic cells (red) is shown on top right. Confocal images depicting peri-LC^GABA^ soma with eYFP (green), Th (white), and Nissl (blue) (bottom) are shown on the bottom. Scale bars are 100 μm, except for inset (20 μm). AP and DV distances in mm from bregma. **(b)** Schematic of whole-cell patch clamp electrophysiology recordings of optically-evoked inhibitory postsynaptic currents (IPSCs). **(c)** Representative optically evoked IPSC is shown (53.8 ± 7.3 pA, n=18) with a paired pulse ratio of 1.1 at a 50 ms interpulse interval (n=16). Blue dashes represent individual pulses of 1 ms at 470 nm 1mW/mm^2^ illumination. **(d)** (Left) Representative recording shows rescue of tetrodotoxin (TTX, 1 μM)-sensitive light-evoked IPSCs by 4-aminopyridine (4-AP, 500μM). (Right) Average IPSCs during 5 min baseline (black), 3-8 min post TTX (orange), and 5-10 min post 4-AP (purple). **(e)** IPSC amplitudes in the presence of TTX or TTX + 4-AP normalized to baseline (n=6). **(f)** Representative trace of a single sweep (top) and an overlay of 10 sweeps (bottom) recorded from an LC^NE^ neuron in cell attached mode. Blue hash marks indicate the duration of light stimulation (1 ms pulses at 20 Hz for 1 s). (Right top) Time course of action potential firing rate for an average of 6 cell-attached recordings with a pulse train for 1 s (20 Hz, 1 ms pulses). (Right bottom) Firing rate before (mean=0.77 Hz), during (0.13 Hz), and after (1.45 Hz) photostimulation. Periods of comparison were 500ms before, middle-to-end, and after stimulation. p=0.0266 for pre-stimulation vs. post-stimulation periods by Wilcoxon signed-rank test. **(g)** (Left) Average amplitudes of optically evoked IPSCs normalized to baseline (n=5) during the wash on of DNQX (10 μM) followed by picrotoxin (100 μM). (Right) Representative traces showing PSCs during 5 min of baseline (black), 5-10 min post DNQX (red), and 5-10 min post picrotoxin (green). **(h)** Schematic of fiber photometry recordings in peri-LC^GABA^ neurons. **(i)** Peri-LC^GABA^ neuronal activity increases during 30s exposure to bobcat urine odor. **(j)** Peri-LC^GABA^ neuronal activity increases during 30s exposure to 2-phenylethanol (2-PE) odor. **(k)** Peri-LC^GABA^ neuronal activity increases after a 1s 0.5mA shock preceded by a coterminating 20s tone. **(l)** Peri-LC^GABA^ neuronal activity increases upon center entry in an open field test. Arrowhead corresponds to entry into the center of the field. **(m)** Schematic (left) of novel object assay; peri-LC^GABA^ neuronal activity increases when animal interacts with a novel object (right), but not a familiar (left) object. Arrowhead corresponds to start of interaction with object. **(n)** Peri-LC^GABA^ neuronal activity decreases during food consumption in food-deprived animal. Arrowhead corresponds to initiation of food consumption. **(o)** Summary of peri-LC^GABA^ fiber photometry experiments. Bars represent change in activity before vs. after events. Periods of comparison were 30s for odor experiments, 20s for shock and conditioned sound, 5s for novel/familiar object interaction and open field exploration, and 10s for feeding. n=12 animals for photometry experiments. *p<0.05, **p<0.01, ***p<0.001, ****p<0.0001 using paired two-tailed t-tests and Friedman ANOVA with Dunn’s multiple comparisons test. ns, not significant. Error bars indicate SEM.

Next, we determined whether peri-LC^GABA^ neurons directly modulate LC activity using *ex vivo* patch-clamp electrophysiology. In acute brain slices from vGAT-Cre mice expressing DIO- ChR2-eYFP in peri-LC^GABA^ neurons (**Fig. 1b**), we recorded from LC^NE^ neurons identified visually and confirmed by the presence of an evoked α_2_-adrenergic receptor-mediated inhibitory current (**Extended Data Fig. 1b**), which is a fundamental characteristic of LC^NE^ neurons^22^. Photostimulation of peri-LC^GABA^ neurons and their terminals evoked fast IPSCs (latency 2.0±0.1ms) insensitive to TTX and 4-AP treatment, demonstrating that peri-LC^GABA^ neurons make monosynaptic inhibitory connections onto LC^NE^ neurons (**Fig. 1c-e**). Photostimulation inhibited ∼78% of LC^NE^ neurons examined (**Extended Data Fig. 1c**), with viral transduction confirmed by observation of fluorescence in the sample. Notably, photostimulation did not affect LC^NE^ neurons when ChR2 was expressed laterally to the LC, indicating that the peri-LC^GABA^ neurons medial to the LC form a distinct population that directly synapses onto the LC. Subsequent experiments only included mice where expression was medial to the LC. Interestingly, photostimulating peri- LC^GABA^ neurons at high frequency (20 Hz, 1 s duration) also caused a pause in LC tonic firing, followed by a rapid burst of LC^NE^ activity upon ceasing stimulation (**Fig. 1f**). Rebound activity in LC^NE^ neurons has been observed in previous studies that directly inhibited the LC^23,24^, which might be mediated by α2-receptor activity^25^. Applying DNQX did not affect optically evoked IPSC amplitude in LC^NE^ neurons, and picrotoxin reduced IPSC amplitude by more than half, indicating that the IPSCs are primarily GABA_A_ receptor-mediated (**Fig. 1g**); we hypothesize that glycinergic currents are responsible for the other portion. Together, these results suggest that peri-LC^GABA^ neurons directly inhibit LC^NE^ activity via GABA release to regulate LC^NE^ firing. We also found that peri-LC^GABA^ neurons were tonically active at baseline, indicating a level of basal tonic inhibition onto LC^NE^ neurons (**Extended Data Fig. 1d-f, Extended Data Table 1**).

### Peri-LC^GABA^ neurons respond to salient stimuli and feeding

To investigate whether peri-LC^GABA^ neurons process arousal-related stimuli, we used fiber photometry to monitor neuronal (calcium) activity in peri-LC^GABA^ neurons. We injected AAV-DIO-GCaMP6s and implanted an optical fiber above the peri-LC of vGAT-Cre mice, then used a battery of diverse behavioral tests in the same animals to determine how a variety of arousing stimuli within different modalities and valence affect peri-LC^GABA^ neuron activity (**Fig. 1h**). First, to determine whether peri-LC^GABA^ neurons respond to odorant stimuli, we exposed mice to 30s of various odors in a chamber with lateral flow. Activity of peri-LC^GABA^ neurons significantly increased upon exposure to the aversive odors of predator urine or 2MT, a derivative of predator urine^26^ (**Fig. 1i, Extended Data Fig. 1g**), to appetitive 2-phenylethanol (2-PE; rose oil extract) or the urine of opposite-sex mice^27,28^ (**Fig. 1j, Extended Data Fig. 1h**), and to peppermint odor as a novel stimulus (**Extended Data Fig. 1i**). Air flow without a corresponding odor did not increase peri-LC^GABA^ GCaMP activity (**Extended Data Fig. 1n**). Taken together, these findings demonstrate that the presentation of salient odors, irrespective of valence, increases peri-LC^GABA^ neuronal activity. As salient stimuli generate phasic bursting in LC^NE^ neurons^29^, it is plausible that peri-LC^GABA^ neurons may suppress certain LC^NE^ subpopulations to increase the relative influence of other LC^NE^ subpopulations, or that peri-LC^GABA^ neurons may receive coincident input to act as a key mechanism for gain control onto LC^NE^ neurons while excitatory inputs coordinate phasic bursting^18^. This tuning of excitatory/inhibitory balance has been described in other brain areas, in which distinct subpopulations of neuromodulatory neurons can differentially influence behavior through their influence on local and distal circuit activity^30–32^.

Second, we determined how peri-LC^GABA^ neurons respond to threat using aversive conditioning to a 20s tone that coterminated with a 1s 0.5mA foot shock. While the foot shock caused a robust increase in peri-LC^GABA^ activity (**Fig. 1k**), the conditioned stimulus alone did not elicit an increase in the GCaMP signal. We also measured peri-LC^GABA^ neuronal activity in less hyper-arousing contexts including the elevated-zero maze (EZM) and a brightly illuminated open field test (OFT). We found that peri-LC^GABA^ neurons increased activity when animals entered the anxiogenic open arm of the EZM or the bright center of the open field, suggesting that these neurons may be engaged during exploration of threatening contexts (**Fig. 1l and Extended Data Fig. 1j-k**). These experiments, however, do not distinguish between bulk versus selective activation of peri-LC^GABA^ neurons. We hypothesize that only a subset of peri-LC neurons are activated so as to suppress selected LC^NE^ subpopulations, while other LC^NE^ neurons undergo the increase in tonic activity associated with anxiety-like behavior^15^.

Next, to determine whether peri-LC^GABA^ neurons respond to novel salient stimuli, as previously established for LC^NE^ neurons^33,34^, we performed novel-object testing as described^20^. In this experiment, peri-LC^GABA^ neurons increased activity during an animal’s interaction with a novel object, but did not change their activity with a familiar object (**Fig. 1m**). This result indicates that the peri-LC^GABA^ neurons respond to novel information, and not to the animal’s conditioned memory of the stimulus^35^. The increased peri-LC^GABA^ activity was surprising, but could represent the activation of a subset of these neurons which inactivate a subset of the LC^NE^ neurons or adjust the excitatory/inhibitory balance to facilitate coordination of phasic bursting of LC^NE^ neurons, similar to the mechanism proposed for local GABAergic control of dopamine activity in the ventral tegmental area^32,36,37^, as well as in cortical projections from the nucleus basalis^31^.

LC^NE^ neurons and a glutamatergic peri-LC neuronal population have been recently implicated in feeding^38^; therefore we sought to determine whether feeding behavior impacts peri- LC^GABA^ activity. In food-deprived animals, peri-LC^GABA^ activity strongly decreased during chow consumption and returned to baseline upon the cessation of consumption (**Fig. 1n-o**). This pattern of activity was also observed when food-deprived animals attempted to eat a false food pellet, which caused a decrease in activity that increased after stopping (**Extended Data Fig. 1l**). When sated mice consumed a highly palatable food item, no significant change in peri-LC^GABA^ activity was observed, though these neurons increased activity when mice ceased an eating bout (**Extended Data Fig. 1m**). This modulation may be associated with switching brain states into and out of a vigilant state^39^. These data indicate that peri-LC^GABA^ neurons are recruited during food consumption behavior in a deprived state. Taken together, these behavioral tests and recordings indicate that peri-LC^GABA^ neurons respond to salient extrinsic arousing stimuli as well as to learned objects and food based on an animal’s homeostatic needs.

### Selective control of peri-LC^GABA^ neuron activity alters arousal states, feeding, and pain responsivity

Because peri-LC^GABA^ stimulation decreases LC activity (**Fig. 1f**), we determined whether physiological and behavioral correlates of arousal and LC^NE^ activity^15,40^ are reduced with peri- LC^GABA^ stimulation. We injected AAV5-EF1a-DIO-hChR2-(H134R)-eYFP and implanted optical fibers bilaterally in the peri-LC of vGAT-Cre mice (**Fig. 2a**). As changes in pupil diameter positively correlate with changes in LC^NE^ activity and global arousal level^18,41^, we measured pupil diameter subsequent to peri-LC^GABA^ photostimulation in anesthetized mice (**Extended Data Fig. 2a**). Photostimulation at 20Hz for 10s decreased pupil diameter in anesthetized mice, consistent with LC^NE^ inhibition (**Fig. 2b**). 20Hz stimulation also caused a decrease in locomotor activity in awake mice in a real-time place preference assay (**Fig. 2c** and **Extended Data Fig. 2b**). Similarly, peri-LC^GABA^ stimulation did not directly modify anxiogenic behaviors in the EZM, open field, and light-dark box, but did decrease the mean velocity of animal locomotion (**Extended Data Fig. 2c-e**). Moreover, stimulating peri-LC^GABA^ neurons did not impair aversive memory acquisition when photostimulation was delivered during tone-shock pairing, nor did it impact novel object interaction or food consumption (**Extended Data Fig. 2f-h**). Peri-LC^GABA^ stimulation was also not positively reinforcing when animals were allowed to nosepoke for photostimulation (**Extended Data Fig. 2i**). Together, these findings suggest that peri-LC^GABA^ neurons can modulate sympathetic arousal and locomotor activity, yet the disparity between photometry and activation results suggest that a more nuanced regulation underlies peri-LC^GABA^ function. Moreover, the suppression of tonic LC^NE^ activity may not eliminate phasic burst firing due to the recently identified robust LC^NE^ excitatory inputs^42^.

**Fig. 2.**
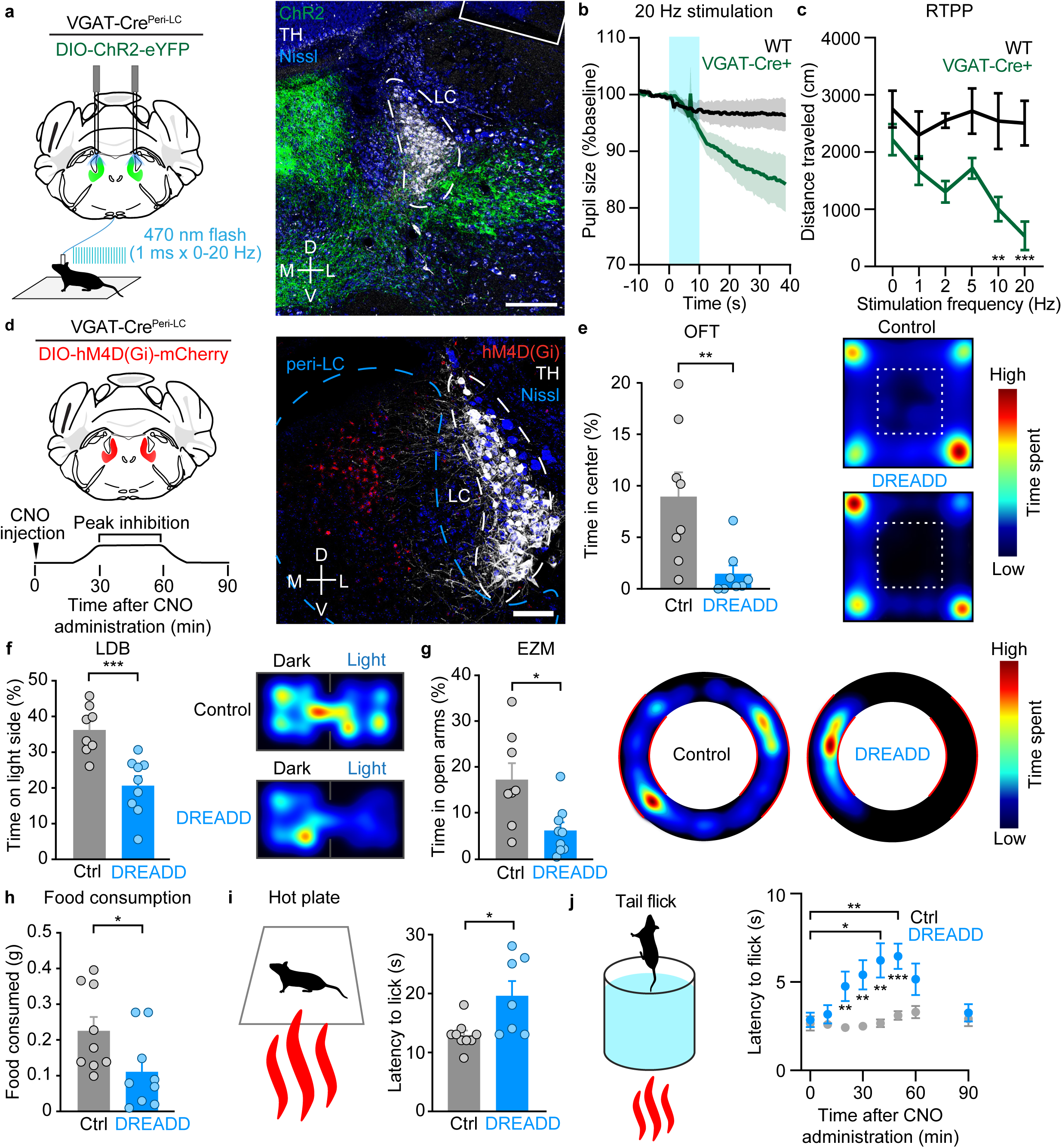
Manipulation of peri-LC GABAergic neurons alters diverse behavioral states. **(a)** (Left) Schematic of viral injection and optogenetic simulation of peri-LC^GABA^ neurons. (Right) Image shows ChR2-eYFP (green), tyrosine hydroxylase (TH, white), and Nissl (blue) expressed in the LC and peri-LC region. Scale bar is 100 µm. **(b)** Graph shows pupil size in response to 10s of 20Hz stimulation (2mW, 5ms pulse width) in wildtype control or vGAT-Cre mice starting at 10s. **(c)** Graph shows distance traveled in real-time place preference assay with varying photostimulation frequencies 0-20 Hz (2mW, 5ms pulse width). **(d)** (Left) Schematic of viral injection and chemogenetic inhibition of peri-LC^GABA^ neurons. (Right) Image shows hM4D(Gi)-mCherry (red), TH (white), and Nissl (blue) in the LC and peri-LC region. Scale bar is 100 µm. **(e-j)** Graphs compare chemogenetic inhibition (1 mg/kg CNO) of peri-LC^GABA^ neurons in vGAT-Cre mice versus non-expressing wild-type controls in **(e)** time spent in the center of an open field, **(f)** time spent on the light side of a light-dark box test (LDB), **(g)** time spent in the open arms of an elevated-zero maze (EZM), **(h)** food consumed in a 20 min period, **(i)** latency to paw lick on a 55°C hot plate, and **(j)** latency to tail flick in 54°C water. Corresponding location heat maps are shown for OFT, LDB, and EZM. *p<0.05, **p<0.01, ***p<0.001 by unpaired two-tailed *t*-test (E-I) or two-way ANOVA (C and J). Error bars indicate SEM.

As peri-LC^GABA^ neurons tonically inhibit the LC at baseline (**Extended Data Table 1**), we hypothesized that inhibiting these neurons would release the LC^NE^ population from inhibitory restraint, in turn increasing anxiogenic and hyperarousal states^18,43^. We injected AAV5-EF1a-DIO- hM4D(Gi), an inhibitory DREADD, into the peri-LC of vGAT-Cre mice (**Fig. 2d**) and measured the behavioral impact of inhibiting peri-LC^GABA^ neurons with 1 mg/kg CNO. Control mice were wild-type C57BL/6J that were injected with the identical virus and received the same dose of CNO. This dose of CNO was used because administering 5 mg/kg CNO to DREADD-expressing mice caused behavioral arrest (**Extended Data Fig. 2j-k**), characterized by a rigid posture and near-zero directed movement, as previously observed when LC^NE^ neurons are hyperactivated^44^. As predicted, selective inhibition of peri-LC^GABA^ increased anxiogenic behavior in the open field (**Fig. 2e** and **Extended Data Fig. 2l**), light-dark box (**Fig. 2f**), and EZM tests (**Fig. 2g** and **Extended Data Fig. 2m**).

A recent study showed that LC activity decreases during food consumption and that LC^NE^ activation attenuates feeding^45^; consistent with this finding, we found that DREADD-mediated inhibition of peri-LC^GABA^ neurons significantly decreased food consumption (**Fig. 2h**). LC activity has differential effects on pain processing based on projection target, wherein activation of LC projections to the spinal cord reduces nociception and hypersensitivity while the activation of projections to the PFC and BLA exacerbate the anxiety and aversion associated with pain ^8,46^. We therefore tested whether inhibiting peri-LC^GABA^ neurons could alter nocifensive behavioral responses. DREADD-expressing animals injected with CNO (1 mg/kg), exposed to a noxious thermal stimulus in the hot plate (**Fig. 2i**) and tail-flick (**Fig. 2j**) assays, indeed showed an increased latency for a nocifensive response relative to non-expressing controls, consistent with an analgesic effect.

Together, these results indicate that peri-LC^GABA^ neurons can bidirectionally modulate global arousal level, affective responses, and nocifensive behavior. Interestingly, both bulk activation and inhibition of peri-LC^GABA^ neurons caused similar effects on behavior, possibly through differential effects on local circuits and distant networks, suggesting that the level of peri-LC inhibition may be more finely regulated. Moreover, the differences we observed between our circuit manipulation and fiber photometry experiments further support the hypothesis that differential activation of peri-LC^GABA^ subpopulations may be a crucial aspect of their function. Together these findings led us to hypothesize that the peri-LC^GABA^ system may be a polymorphic population of neurons with heterogeneous functional roles which influence multiple types of arousal-like and avoidance behaviors.

### LC and peri-LC neurons are composed of diverse molecular subpopulations

Having observed diverse and sometimes counter-intuitive LC and peri-LC^GABA^ neuronal activity responses during different behavioral paradigms, we next posited that molecular heterogeneity of LC and peri-LC^GABA^ neuronal subpopulations may contribute to these functional variations. To better understand the cellular diversity within the LC and peri-LC, we used single-nucleus RNA sequencing (snRNAseq) in brain sections containing both the LC and peri-LC from adult wild-type male or female mice and processed them by droplet-based nuclear mRNA partitioning and sequencing (**Fig. 3a**, see methods and **Supplementary Table S1** for details)^47^. After filtering low-quality cells and doublets, we identified 30,818 total cells (19,245 male and 11,573 female) across 3 samples, each pooled from 5-6 animals, and performed graph-based weighted nearest neighbor clustering analysis. We classified 15,050 cells (48.8%) as neurons, based on their expression of the canonical neuronal markers *Stmn2* and *Thy1* (**Fig. 3b-c; Extended Data Fig. 3a-g**)^47^. We excluded VGLUT1 neurons in our samples because they were of low-quality (**Extended Data Fig. 3e-g**), were detected in only a single sample, and *Slc17a7* (VGLUT1) was not expressed in the medial peri-LC zone in fluorescent *in situ* hybridization (FISH) (**Extended Data Fig. 3k**).

**Fig. 3.**
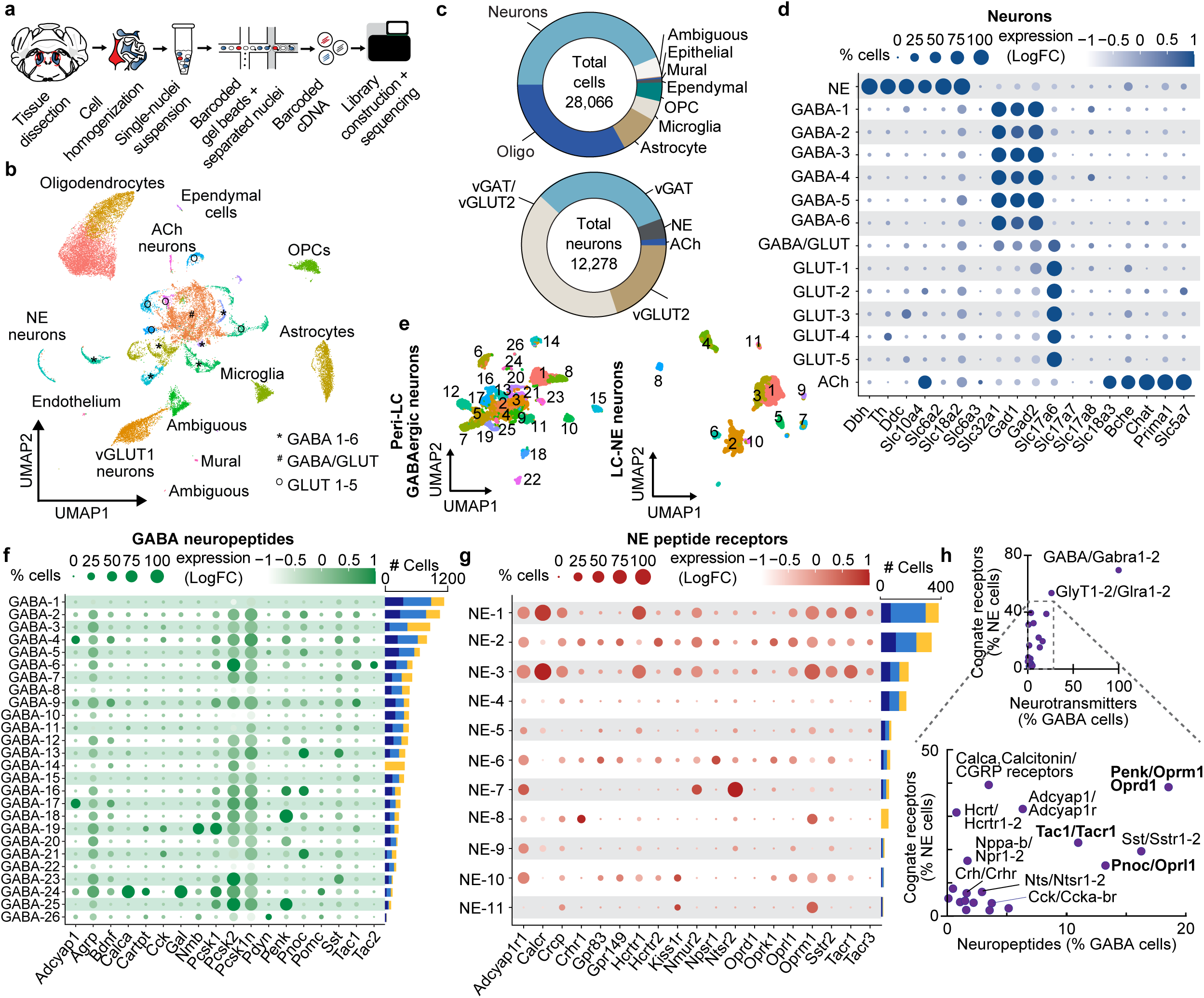
Isolating the locus coeruleus and peri-LC transcriptome. **(a)** Schematic shows the steps in the single-nucleus RNA sequencing (snRNAseq) experiments. **(b)** UMAP visualization based on transcriptional profile is shown for n=30,818 cells pooled from 3 different samples. **(c)** Distribution of cell types (top, n=28,066) and neuronal classes (bottom, n=12,278) in snRNAseq sample after vGLUT1 neurons were removed. **(d)** Dot plots of n=12,278 neurons are shown, with neuronal clusters on the y-axis and gene transcripts on the x-axis. Size of circles corresponds to percent of cells in the cluster expressing a specific transcript, while color intensity corresponds to the relative expression level of the transcript. Plots correspond to transcripts related to neurotransmission. **(e)** UMAP visualizations of reclustered neurons using *Slc32a1* (vGAT), *Gad1*, or *Gad2* expression for the peri-LC inhibitory neurons (left, n=11,182); and *Dbh* or *Th* expression for the LC^NE^ neurons (right, n=1,324). **(f)** Dot plots of the subset of neurons that express *Slc32a1* (vGAT), *Gad1*, or *Gad2* (n=11,182) are shown, with the GABAergic subpopulations on the y-axis and neuropeptide genes on the x-axis. Circle size corresponds to percent of cells in the cluster expressing a specific transcript, while color intensity corresponds to its relative expression. Cell number in each cluster is graphed to the right, where the yellow represents the contribution from the female sample and blue represents the two male samples. **(g)** Dot plots of the subset of neurons that express *Dbh* or *Th* (n=1,324) are shown, with LC^NE^ subpopulations on the y-axis and neuropeptide receptor genes on the x-axis. Circle size corresponds to percent of cells in the cluster expressing the specific transcript, while color intensity corresponds to its relative expression. Cell number in each cluster is graphed to the right, where the yellow represents the contribution from the female sample and blue represents the two male samples. **(h)** Graphs show the distribution of neurotransmitter/neuropeptide gene expression in GABAergic neurons on the x-axis and the gene expression of their cognate receptors in LC^NE^ neurons on the y-axis.

Interestingly, LC and peri-LC neurons formed clusters based on the glutamate, GABA, and the monoamine neurotransmitters they release (**Fig. 3d; Extended Data Fig. 3h**). The LC^NE^ cluster was highly enriched in genes involved in NE synthesis and transport, including *Dbh*, *Th*, *Ddc*, *Slc10a4*, *Slc6a2* (NET), and *Slc18a2* (VMAT2), but low in *Slc6a3* (DAT), the gene involved in dopamine transport, suggesting that these are predominantly LC^NE^ neurons and not dopamine neurons^48^. GABAergic neurons, on the other hand, were separated into 7 clusters that displayed high expression of *Slc32a1* (VGAT), *Gad1*, and *Gad2*, with one cluster co-expressing high levels of *Slc17a6* (VGLUT2). Five additional classes of VGLUT2 neurons were identified, but no cluster highly expressed *Slc17a7* (VGLUT1) or *Slc17a8* (VGLUT3). The last cluster of neurons expressed *Slc10a4*, *Slc18a3*, *Bche*, *Chat*, *Prima1*, and *Slc5a7*, which are typically characteristic of cholinergic (ACh) neurons^49^.

We identified LC and peri-LC neurons which expressed several neuropeptides together with their cognate G-protein coupled receptors (GPCRs), which are likely candidates for modulatory tuning of diversity in LC function (**Extended Data Fig. 3i-j**). LC^NE^ neurons were highly enriched in *Calca* (calcitonin/CGRP), *Cartpt*, and *Gal* (galanin), as well as *Calcr* (calcitonin receptor). Peri-LC^GABA^ neurons, on the other hand, expressed high levels of *Penk*, *Pnoc*, and *Tac1*. Peri-LC VGLUT2 neurons expressed high levels of *Adcyap1*, which encodes PACAP, with subpopulations also expressing *Cck* or *Bdnf*. *Nps* was expressed across all populations.

The LC has long been associated with opioid neuropeptide and receptor signaling in stress, pain, and analgesia^50,51^. *Pdyn* (preprodynorphin) expression was low, except within one glutamatergic population (GLUT-4), but *Oprk1* (kappa opioid receptor) was primarily expressed in glutamatergic neurons, consistent with previous studies showing that dynorphin signaling likely acts on presynaptic terminals in the LC region^52,53^. *Oprm1* (μ opioid receptor) expression was highest in LC^NE^ neurons, consistent with previous studies^54,55^; expression was also common across many non-noradrenergic groups. *Hcrtr1* and *Hcrtr2*, the two orexin receptors, were expressed on different cell populations, indicating potentially differential roles in regulating arousal, motivation, and opioid abuse-related behaviors^56,57^.

Additional analyses were performed to characterize the transcriptional diversity in physiology, structure, and disease-associated genes within the LC and peri-LC. We found differential expression in neuropeptide signaling, neurotransmitter reuptake, endocannabinoid signaling, hormone receptors, LC developmental markers^58–60^, and genes involved in cell morphology and adhesion (**Extended Data Fig. 4a-c**). Differential expression of gap junction proteins (**Extended Data Fig. 4b**) may allow LC ensembles to form by electrotonic coupling and may explain sub-millisecond LC synchrony^19,61^. Differences in the expression of genes associated with psychiatric diseases, such as *Maoa*, *Ptger3*, and *Peg10*^62^ (**Extended Data Fig. 4d**), and with dementia, such as genes involved in amyloid and tau interactions (**Supplementary Table S2**) were also found in our sample. We also report heterogenous expression of ion channels, genes associated with neurotransmitter loading and release, and GPCRs and genes associated with GPCR regulation (**Supplementary Table S2**).

To investigate whether heterogeneity may exist within LC^NE^ and peri-LC^GABA^ neurons, we next sought to identify subpopulations by performing conservative reclustering only in neurons expressing *Dbh* or *Th* (**Fig. 3e-f; Extended Data Fig. 3l**); or expressing *Slc32a1*, *Gad1*, or *Gad2* (**Fig. 3e,g; Extended Data Fig. 3m**). Reclustering parameters were determined by using ChooseR subsampling to identify maximally robust clustering resolution^63^, which was 1.2 for GABAergic neurons and 0.8 for NE neurons **(Extended Data Fig. 3l-m**). Given other recent reports describing sexual dimorphism in LC structure and function^62,64,65^, we also identified sexually dimorphic genes in all cells, the LC^NE^ subset, and the peri-LC^GABA^ subset (**Supplementary Table S3**). We found surprisingly few sex differences in gene expression, none of which would be expected to have significant functional effects. Since a previous study of LC^NE^ gene translation uncovered much more significant sexual dimorphism^62^, we suspect that the major contributor to sexually dimorphic gene expression in the LC derives from post-translational regulation, which may be connected to the increased number of long noncoding RNAs in females or to robust brain state changes during periods of stress (**Supplementary Table S3**). Indeed, all clusters were made up of cells from all samples, with the exception of one GABAergic and one NE cluster (GABA-14 and NE-8) (**Fig. 3f-g**), which were exclusively female, and likely grouped by the expression of these noncoding RNAs, as they show no other functional distinctions.

Subpopulations of LC^NE^ neurons defined by co-expression of glutamatergic markers and GABAergic markers may indicate a role in lateral inhibition or excitation within the LC^NE^ cells. Striking differences in expression are found in the gap junction proteins (**Extended Data Fig. 5c**), which may underlie some of the previously reported electrotonic coupling between LC^NE^ ensembles^19,66^. LC^NE^ neurons also differ in their receptors for neurotransmitters and neuromodulators, including α_2_-adrenergic receptors (**Fig. 3g; Supplementary Table S2**) and in ion channel expression (**Supplementary Table S2**), which may facilitate the heterogeneous, intrinsic, electrophysiological differences in LC neurons^6,67,68^, rebound activity^25^, and responses to synaptic inputs. We also found that peri-LC^GABA^ subpopulations exhibit key differences (**Fig. 3f**), including differential expressions of *Penk*, *Pnoc*, and *Tac1* neuropeptides. Interestingly, GABA- 3, GABA-13, and GABA-21 neurons co-express *Pnoc* and *Slc6a5* (glycine transporter) (**Extended Data Fig. 6b**), suggesting that peri-LC^Pnoc^ neurons may be glycinergic. Taken together, the single-nucleus transcriptional profiles reveal distinct and molecularly heterogenous subpopulations of LC and peri-LC neurons.

### LC and peri-LC RNA transcripts are spatially distributed in anatomically discrete locations

We next corroborated and extended our snRNAseq findings by determining the spatial distribution of RNA transcripts using an recently developed approach called Pixel-seq, in which a gel covered by high-density DNA polymerase colonies (polonies) allows capture of tissue mRNA followed by sequencing of RNA transcripts along with spatially-identified barcodes^69^ (**Fig. 4a**). Matching the barcodes to their position in the gel allows for the segmentation of highly spatially resolved transcripts into individual cells. Tissue containing the LC and peri-LC was dissected from 10μm-thick sections, its RNA hybridized to the barcoded polony gel, and the cDNA then amplified and sequenced. Coronal sections (6 female, 5 male) were collected of the LC and peri-LC along the anterior-posterior axis, with a representative unique molecular identifier spatial density map of the region shown in **Fig. 4b**, and total for each section shown in **Extended Data Fig. 7e**. Read saturation exceeded >70% for both male and females, with similar gene abundance; and ∼90% of the mapped transcripts were from protein-coding genes (**Extended Data Fig. 7b-d**). Leveraging cell type-enriched markers found in our snRNAseq experiments, we detected ∼59,000 cells (**Extended Data Fig. 7a**) and clustered them into neurons and non-neuronal subtypes (**Fig. 4b**). Among the neurons, we further identified subpopulations, which when spatially mapped, showed a dense core of noradrenergic neurons surrounded by a group of GABAergic neurons (**Fig. 4c-d**), consistent with our anatomical studies we report in **Fig. 1a**. Established noradrenergic markers (*Dbh*, *Th, Slc18a2* and *Gal*) were densely expressed in the LC, in contrast to the various neuropeptides identified by snRNAseq (*Calca*, Hctr1, *Npy*, *Nps*, *Penk*, *Pnoc*, *Sst*, and *Tac1*), which were broadly distributed throughout the peri-LC (**Fig. 3c-d, Extended Data Fig. 7f**). Interestingly, *Penk*, *Sst*, and *Nps* appeared to express more highly in the posterior peri-LC (**Fig. 3e**), with *Sst* showing a distinct cluster ventromedial to the LC.

**Fig. 4.**
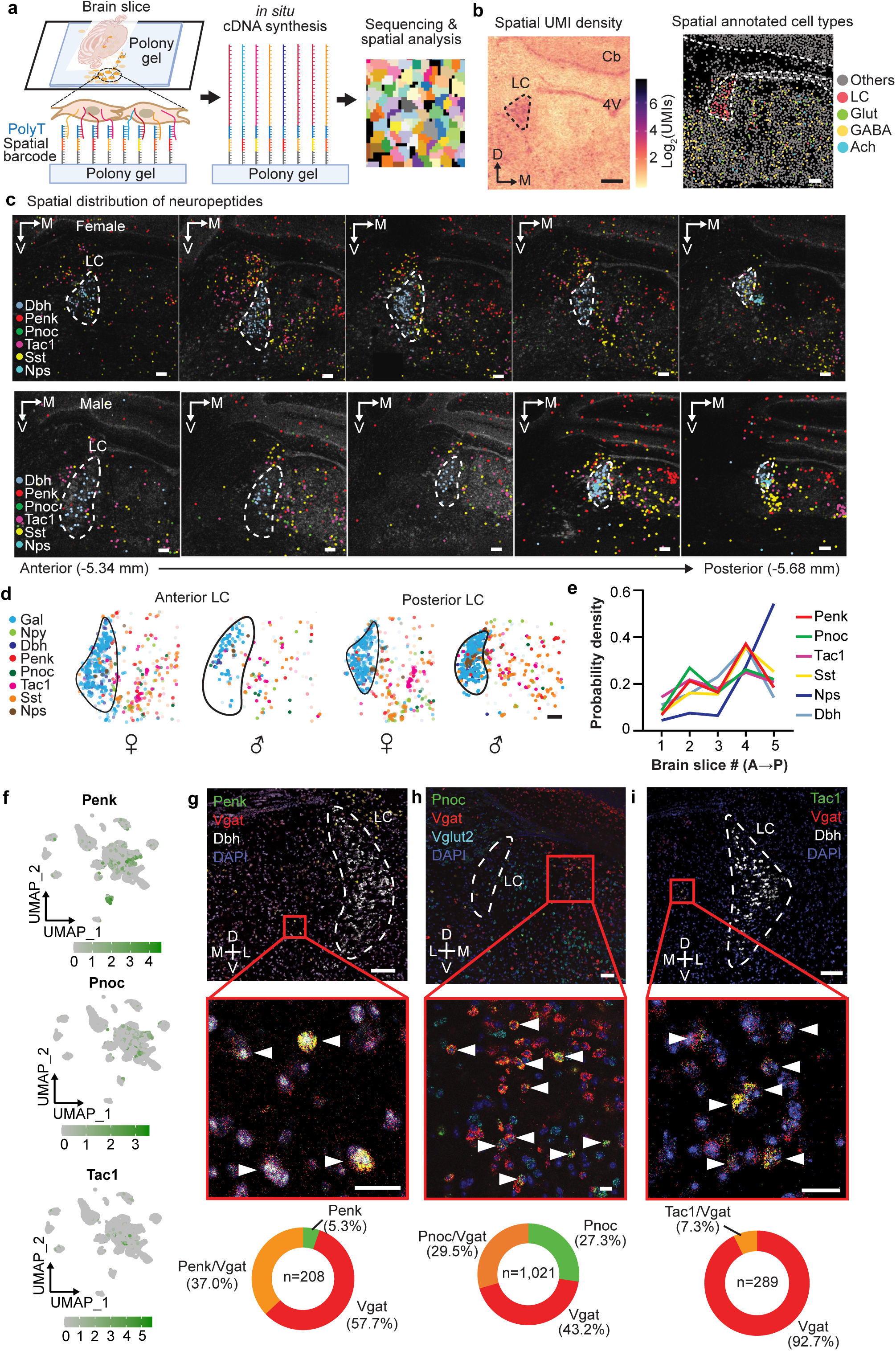
Spatial transcriptomics using PIXEL-seq of the LC and peri-LC region. **(a)** Schematic depicts spatial transcriptomics via Pixel-seq. Barcoded DNA templates were grafted onto polyacrylamide gel and the spatial coordinates of barcodes were determined by Illumina sequencing by synthesis method. The right plot shows the spatial distribution of unique molecular identifier (UMI) density per cluster (1 pixel = 0.325 x 0.325 µm^2^). **(b)** Left, a spatial UMI density image of a representative fresh tissue section containing the LC and neighboring regions. 4V, 4^th^ ventricle. CTX, cerebellum. Scale bar 200 µm. Right, single-nucleus RNA sequencing-guided annotation of segmented pseudo-cells is shown for neuronal and non-neuronal subpopulations. Scale bar is 100µm. **(c)** Spatial expression maps of *Dbh, Penk, Pnoc, Tac1, Sst, Nps* of the LC region along the anterior-posterior axis are shown for a female (top) and male (bottom) mouse. Scale bar 100 µm. **(d)** Summary comparisons of the spatial expressions of *Gal, Npy, Dbh, Penk Pnoc, Tac1, Sst, Nps* in the anterior and posterior LC in male and female mice. Each pseudo-cell has been enlarged 1.5 times to aid with visualization. Scale bar is 100µm. **(e)** Probability density of the expressions of various neuropeptide and *Dbh* transcripts along the anterior-posterior axis (Bregma -5.34 mm to -5.68 mm) in the LC region. **(f)** Feature plots show the peri-LC inhibitory neuron clusters expressing Penk (top), Pnoc (middle), and Tac1 (bottom). **(g-i)** Representative images for *in situ* hybridization of vGAT (*Slc32a1*) (**g-i**), *Dbh* (**g,i**), and vGLUT2 (*Slc17a6*) (**h**) with (**g**) *Penk*, (**h**) *Pnoc*, **(i)** *Tac1* mRNA. Scale bars are 100μm (top) and 20μm (bottom). The respective quantification of the relevant *in situ* cell populations within the peri-LC are shown in the bottom row.

To better appreciate the functional significance of genetically distinct peri-LC clusters in behavior, we selected three peptidergic subpopulations to investigate further: pENK, Pnoc, and Tac1 (**Fig. 4f**). This was based on their high gene expression in the peri-LC^GABA^ population and the high gene expression of their cognate receptors within the LC^NE^ clusters (**Fig. 3h; Extended Data Fig. 3n-o**). We validated the peri-LC expression of these peptide mRNAs using FISH (**Fig. 4g**), and then examined the responses of these subpopulations during behavioral studies using photometry.

### Peri-LC neuropeptidergic subpopulations have differential activation patterns and effects on the LC

We hypothesized that the peptidergic peri-LC subpopulations could have diverse functions, as several prior reports have suggested that different neuropeptide GPCRs can regulate LC^NE^ function^15,70–74^ in various ways. Based on isolation of highly expressed peptide genes in the peri- LC^GABA^ population and the presence of cognate GPCR genes within the LC^NE^ population (**Fig. 3h**), we investigated peptidergic neurons expressing pENK (13.4% of peri-LC population), Pnoc (8.0% of peri-LC population), and Tac1 (10.2% of peri-LC population), which also have limited overlap in expression within the peri-LC (**Fig. 4f; Extended Data Fig. 3p-q**). We injected AAV-DIO-GCaMP6s and implanted an optical fiber in the peri-LC of vGAT-Cre, pENK-Cre, Pnoc-Cre, and Tac1-Cre mice to determine how different stimuli impact the activity of these peri-LC subpopulations (**Fig. 5a**-**b**, **Extended Data Fig. 8a**).

**Fig. 5.**
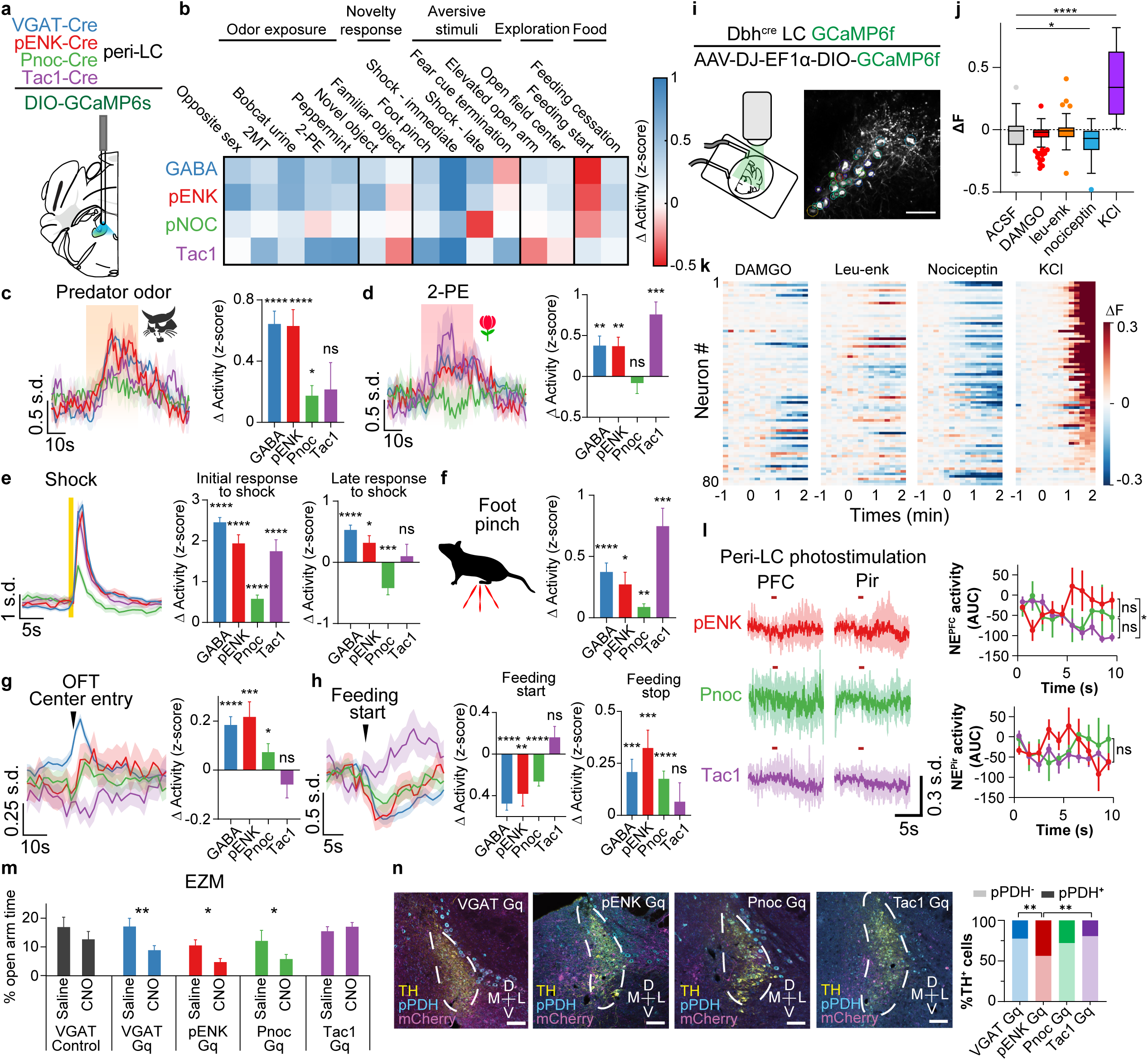
Functional heterogeneity of peri-LC neuronal subpopulations. **(a)** Cre-dependent GCaMP6s was expressed in the peri-LC of vGAT-Cre, pENK-Cre, Pnoc-Cre, or Tac1-Cre mice and recorded with fiber photometry. **(b)** Heat map summary of peri-LC fiber photometry experiments. **(c-h)** Photometry traces and their quantification for the peri-LC neuronal subpopulations (vGAT, pENK, Pnoc, Tac1) are shown for responses to 30s exposure to bobcat urine odor **(c)**, to 30s exposure to 2-phenylethanol (2-PE) odor **(d)**, to early (left, 0-5 s after shock) and late (right, 5-20s after shock) responses to a 1s 0.5mA footshock **(e)**, to a foot pinch **(f),** to entering the center of an open field **(g)**, and to the start and end of a food consumption bout **(h)**. Bars represent change in activity before vs. after events. Periods of comparison were 30s for odor experiments and foot pinch, 5s for novel/familiar object interaction and open field exploration, and 10s for feeding. Early shock comparison period was defined as 5s before and after shock initiation; late shock comparison period was defined as 5-20s after shock and 5-20s before shock. n=12 animals for GABA photometry, 6 for pENK, 5 for Pnoc, and 4 for Tac1. *p<0.05, **p<0.01, ***p<0.001, ****p<0.0001 by paired two-tailed *t*-test. Error bars indicate SEM. **(i)** Left schematic shows 2-photon setup for slice imaging and solution exchanges. Right shows an example brain slice with GCaMP6f-expressing LC^NE^ neurons of Dbh-Cre mice. Scale bar 100 µm. **(j)** Box plots show fluorescent changes of GCaMP6f-expressing LC^NE^ cells after perfusing with either ACSF (control), 1-4 µM DAMGO, 1 µM Leu-enk, 1 µM nociceptin, or 30mM KCl. **p<0.01. ****p<0.0001 by two-tailed *t*-test after a significant ANOVA test. Error bars indicate SEM. **(k)** Heat plots of individual neurons (n=80 from 5 slices) show responses to DAMGO, Leu-enk, nociceptin, and KCl. **(l)** Left, NE activity measured by photometry traces of GRAB-NE2m sensor, aligned to peri-LC photostimulation (30Hz 500ms every 20s repeated 50 times), averaged across all events and across mice for pENK-Cre (N=6), Pnoc-Cre (N=5), and Tac1-Cre (N=8) mice in the PFC (top) and Pir (bottom). Right, plots show 1-s binning of NE changes in response to photostimulation at time 0. *p<0.05 by Spearman correlation. Error bars indicate SEM. **(m)** Change in the percent time spent in center of an elevated zero maze calculated as the full 20 min of assay ∼30 min after saline or 1 mg/kg CNO treatment. Comparisons are shown for VGAT fluorophore control (N=7), VGAT Gq (N=7), Penk Gq (N=10), Pnoc Gq (N=9), and Tac1 Gq (N=13) mice. *p<0.05, **p<0.01 by 2-way ANOVA. Error bars indicate SEM. **(n)** Immunostains for phospho-pyruvate dehydrogenase (pPDH) as marker for neuronal silencing after activating Gq-DREADD expressed in select peri-LC populations. Right, bar graph shows the corresponding quantification of the % pPDH positivity in TH^+^ locus coeruleus neurons after activating the Gq DREADD. **p<0.01 by chi-square test of homogeneity with Bonferroni correction.

Exposure to a predator odor significantly increased the activity of peri-LC GABAergic, pENK, and Pnoc populations, but not of Tac1 neurons (**Fig. 5c**). The same three populations also increased activity when presented with the odor of opposite-sex mice (**Extended Data Fig. 8b**), demonstrating that the valence of the stimulus did not affect the activity of these populations. Curiously, exposure to 2MT (**Extended Data Fig. 8c**), 2-PE (**Fig. 5d, Extended Data Fig. 8m**), and peppermint odor (**Extended Data Fig. 8d**) caused a significant increase in activity in the peri-LC GABAergic, pENK, and Tac1 populations, but not in Pnoc neurons. Control air flow did not affect the activity of any peri-LC population (**Extended Data Fig. 8e**) we tested. These experiments demonstrate that peri-LC subpopulations exhibit diverse physiological responses to a variety of salient odorant stimuli. We also found that peri-LC^GABA^ and peri-LC^pENK^ neurons show increased activity when an animal interacted with a novel object but did not change significantly when exposed to a familiar object (**Extended Data Fig. 8f**). By contrast, peri-LC^Tac1^ neurons increased activity during novel object interactions, but decreased activity during familiar object interactions (**Extended Data Fig. 8f**). Together, these results indicate that peri-LC^GABA^, peri-LC^Pnoc^, peri-LC^Tac1^, and peri-LC^pENK^ populations are likely represent distinct peri-LC nodes which differentially integrate novel stimuli.

We also examined the activity of peri-LC subpopulations during food consumption. On one hand, the GABAergic, pENK, and Pnoc populations decreased activity during chow consumption in food-deprived animals, while they increased activity after finishing consumption (**Fig. 5h**). Tac1 neurons, on the other hand, did not significantly change activity during or immediately after consumption. Additionally, no population that we recorded from changed activity during the consumption of highly palatable food when sated, with only the peri-LC^GABA^ neurons increasing in activity after an eating bout (**Extended Data Fig. 8j**), potentially suggesting that an unidentified subpopulation of peri-LC^GABA^ neurons likely modulates hedonic feeding or may contribute to the change in brain state associated with the termination of feeding in favor of observing or exploring their environment.

We next determined whether activity of the peri-LC neurons differentially respond to hyperarousing stimuli, including threat and anxiogenic contexts, using fear conditioning (**Fig. 5e**), EZM, and open field tests. In aversive conditioning, pre-testing showed no inherent response to the tone (**Extended Data Fig. 8g**) in any of the four peri-LC populations, but the footshock caused a sharp increase in activity in all cell populations. However, after this initial increase, the peri-LC GABAergic and pENK neurons remained elevated in activity, whereas Pnoc neurons decreased activity to below pre-shock baseline (**Fig. 5e**). Interestingly, the next day, none of these peri-LC populations responded to the tone onset; however, peri-LC^GABA^ neurons showed a small but significant decrease in activity at the end of tone presentation, while Tac1 neurons sharply increased activity at tone offset (**Extended Data Fig. 8h**). Another noxious stimulus, a foot pinch, also elicited increased activity in all four populations (**Fig. 5f**). In experiments involving anxiogenic contexts, we found that peri-LC^GABA^ neurons increased activity when animals entered the open arm of an EZM or the center of an open field, whereas pENK and Pnoc populations increased activity only in the center of the open field, but not upon entering the open arm of an EZM (**Fig. 5g** and **Extended Data Fig. 8i**). Peri-LC^Tac1^ neurons, by contrast, did not change activity in the center of an open field, but decreased activity in the open arm of an EZM. Combined, these experiments indicate that peri-LC^pENK^ and peri-LC^Pnoc^ populations are recruited during exploration of an anxiogenic exposed environment, while Tac1 neurons may decrease activity during exposed and elevated exploratory behavior. Peri-LC^GABA^ neuron activity was correlated with movement in both the EZM and OFT, consistent with their role in exploratory behavior, while peri-LC^pENK^ neurons were active during movement in the OFT and peri-LC^Pnoc^ neurons were active during movement in the EZM (**Extended Data Fig. 8k-l**).

To survey how these neuropeptidergic peri-LC inputs may act to modulate the LC, we obtained brain slices expressing GCaMP6f in the LC of Dbh-Cre mice and washed on several neuropeptides including DAMGO, leu-enkephalin, and nociceptin while monitoring LC calcium responses at the single-cell resolution with a 2-photon microscope (**Fig. 5i**). We imaged a total of 80 LC^NE^ cells that increased fluorescence to 30mM KCl, confirming their responsiveness. When all cells were considered, nociceptin exposure significantly decreased neuronal activity across the LC^NE^, but no significant changes were observed in the combined population response to DAMGO, leu-enk, and control ACSF (**Fig. 5j**). Given the literature demonstrating potent mu-opioid mediated hyperpolarization via GIRK activation in LC^NE^ neurons^75,76^, the lack of widespread response to DAMGO and leu-enk was surprising; however, a subset of DAMGO-treated neurons (shown as Tukey outliers) did exhibit decreased activity, consistent with heterogeneity in neuronal responses. Critically, the three different agonists differentially activated or suppressed neuronal activity at the single-cell level (**Fig. 5k**), a heterogeneity reflected in the lack of normality and correlation (**Extended Data Fig. 8n**) in the neuronal responses. Thus, these results suggest that the neuropeptides generated by peri-LC GABA neurons can heterogeneously influence the activity of individual LC^NE^ neurons, which may result from differential expression of peptidergic receptors in LC^NE^ neurons (**Fig. 3g-h**).

We then determined whether the activation of these three peri-LC peptidergic subpopulations may differentially affect LC^NE^ function at the circuit level. The opsin ChrimsonR was selectively expressed in each peri-LC subpopulation using pENK-Cre, Pnoc-Cre, and Tac1- Cre mice, and the recently developed fluorescent NE sensor GRAB_NE2m^77,78^ was expressed in the medial prefrontal cortex (PFC) and the piriform cortex (Pir). Optical fibers were implanted in the peri-LC for stimulation and in the PFC and Pir for measuring NE release dynamics. Although photostimulating the peri-LC pENK, Pnoc, and Tac1 subpopulations at 30Hz for 500ms did not produce any significant differences in Pir NE release, PFC NE release differed significantly in response to activation of the pENK subpopulation compared to the Tac1 subpopulation (**Fig. 5l, Supplementary Table S1**). We also used a tail lift stressor, which is known to robustly increase downstream LC^NE^ activity^79^ (**Extended Data Fig. 8o**), and in each genotype we found equivalent PFC and Pir NE release dynamics following tail lift stress, further indicative of differences between global versus discrete modes of peri-LC activity which may contribute to polymorphic LC^NE^ function^4^. Taken together, these results indicate that peri-LC neuropeptide subpopulations can have differential effects on LC^NE^ system function.

We subsequently determined whether activation of peri-LC subpopulations may differentially impact avoidance behaviors. VGAT-Cre, pENK-Cre, Pnoc-Cre, and Tac1-Cre mice were injected with an activating designer receptor (Gq-DREADD), given either saline or CNO, and then tested in the EZM and OFT. A group of mice expressing only a fluorophore was used as a control. As expected, in the control group, time in the open arms of EZM and in the center of the OFT did not differ (**Fig. 5m, Extended Data Fig. 8q**). Consistent with previous findings showing that LC^NE^ neuron activity is necessary and sufficient for regulation of avoidance and exploration in an aversive environment^43^, DREADD stimulation of peri-LC GABA, pENK, and Pnoc neurons decreased time spent in the open arms of the EZM as compared to the saline condition. Stimulation of Tac1 neurons caused no significant difference in either in the OFT or EZM. These groups of mice did not differ in their distance traveled, except for the VGAT Gq group in the EZM (**Extended Data Fig. 8p**). These results, in combination with our prior results, indicate that the peri-LC subpopulations we identified in this study likely influence LC^NE^ function at the behavioral level, though the regulation of distinct behaviors may be more complex.

Finally, to assess whether exciting the different peri-LC populations would lead to differences in LC^NE^ neuron inhibition, we used an antibody against phospho-pyruvate dehydrogenase (pPDH) to immunostain putatively inhibited neurons^80^. We observed robust pPDH expression in the LC-NE cells of mice in which we activated select peri-LC neurons (**Fig. 5n**) with CNO injection, and further quantification revealed a significantly higher proportion of pPDH^+^ tyrosine hydroxylase (TH^+^) neurons in response to the activation of peri-LC^pENK^ neurons compared to peri-LC^Tac1^ or peri-LC^GABA^ neurons. Together, these convergent results across numerous experiments support the hypothesis that activation of selected peri-LC subpopulations is a critical mechanism by which the LC coordinates differential shifts in excitatory/inhibitory balance with LC^NE^ neurons, subsequently modulating arousal and arousal-related behaviors.

## Discussion

Here we define a heterogenous peri-LC^GABA^ neuronal population whose inhibitory inputs modulate LC^NE^ firing and arousal, avoidance and related exploratory behaviors. We identify molecular diversity within LC and peri-LC and provide evidence which suggests nonuniform spatial organization within this region. While the functional implications of this organization require further study, knowing the spatial expression patterns and diversity in accessible neuronal types will allow many new experimental questions can be addressed.

We then survey the peri-LC^GABA^ and peptidergic neuron type activity during exploratory behavior, feeding, associative learning, spatial and object exploration, and processing of odor and pain stimuli. We report that increasing the activity of peri-LC^GABA^ neurons rapidly but temporarily inhibits tonic activity of LC^NE^ neurons, in addition to inhibiting LC phasic bursting, as described before^81^. Additionally, given the predominantly local projections of peri-LC^GABA^ neurons (**Fig. 1a**), and that most, if not all, LC^NE^ neurons were subject to modulation by peri-LC^GABA^ neurons, peri-LC^GABA^ likely controls LC^NE^-mediated arousal states and responses via direct interactions. Consistent with this idea, we found that activating peri-LC^GABA^ neurons decreased global arousal and exploratory behavior (**Fig. 1f**, **2b**-**c**), whereas inhibiting these neurons increased avoidance behaviors (**Fig. 2d-g**), consistent with disinhibition of LC activity^82^. To explore any potential functional diversity of these populations, we defined the endogenous activity of peri-LC^GABA^ neurons in multiple behavioral domains that LC^NE^ system has been implicated in controlling. These included arousing, aversive and appetitive stimulus processing (**Fig. 1i-j, Extended Data Fig. 1g-i**), sexual behavior (**Extended Data Fig. 1a**), feeding (**Extended Data Fig. 1l**), nocifensive response (**Fig. 1k, 4k**) and novel object processing (**Fig. 1m**). The peri-LC^GABA^ response was present across all modalities tested except for the auditory tone (**Extended Data Fig. 8g-h**), which was surprising in light of previous findings identifying short-lived (<100 ms) tone onset responses in a subset of peri-LC^GABA^ neurons using electrophysiology^18^. We suspect that the tone onset response may have been washed out by opposing signals in the bulk photometry recordings, or that the temporal precision of GCaMP6s is insufficient to detect these changes. Due to the activation of peri-LC^GABA^ by similar inputs that activate LC^NE^ neurons, as well as significant overlap in regions that project to the peri-LC^GABA^ and LC^NE^ populations^18^, it is likely that common inputs to the two populations influence the LC^NE^ neurons by adjusting the excitatory/inhibitory balance, as has been identified in other neuromodulatory centers^30,31^.

Two complimentary transcriptomic approaches revealed significant molecular and spatial diversity in peri-LC^GABA^ neurons which we report translates to functional diversity in a variety of settings. In particular, peri-LC^GABA^ neurons expressing a wide repertoire of neuropeptides (**Fig. 3f**) surround the LC^NE^ dendritic field as revealed by our Pixel-seq spatial transcriptomic results (**Extended Data Fig. 7g**). As differing groups of LC neurons also express diverse sets of cognate receptors (**Fig. 3g**, **Extended Data Fig. 5a**), the combinations of neuropeptide-receptor interactions between the peri-LC and LC are likely to generate polymodal LC ensembles that can flexibly alter their dynamics and E/I balance depending in response to different internal states and behavioral needs. For instance, given that peri-LC^GABA^ activity decreased during food consumption (**Fig. 1n**), we hypothesized that inhibiting these neurons would increase feeding, but chemogenetic inhibition of these neurons instead decreased feeding (**Fig. 2h**), likely by disinhibiting LC^NE^ projections to the lateral hypothalamus involved in feeding^45^. This seemingly paradoxical finding may be due to chemogenetic inhibition affecting a less specific population of neurons than those involved in natural feeding behavior, which could engage vastly different LC^NE^ ensembles and projections (compare ^83^ vs. ^45^). We also found that inhibiting peri-LC^GABA^ neurons promoted analgesia (**Fig. 2i**-**j**), likely by disinhibiting LC^NE^ projections to the dorsal horn of the spinal cord, which act to inhibit noxious afferents^84,85^, rather than LC^NE^ projections to the BLA^46^ and PFC^8^, which contribute to negative affect in pain response; however, a direct test of this concept requires further investigation. Furthermore, although novel and salient stimuli were generally associated with increased peri-LC GABAergic and peptidergic activity (**Fig. 5b,e**), Tac1 neurons decreased activity during exploration in contrast to peri-LC^pENK^, peri-LC^Pnoc^, and the peri- LC^GABA^ population as a whole (**Fig. 5g, Extended Data Fig. 8i**). While peri-LC^Tac1^ neurons were the most functionally distinct from other cell types in behavior, they also have the highest overlap with non-GABAergic neurons in our sequencing results (**Extended Data Fig. 3p-q**); however, this likely includes neurons lateral to the LC (**Fig. 5c**), which were excluded from our photometry datasets (**Extended Data Fig. 8a**). Importantly, the number of cells within these subpopulations and their distribution throughout the peri-LC region varied, complicating direct comparison between their fiber photometry signals; single-cell 1-photon or 2-photon imaging studies are likely to facilitate more direct comparisons. The molecular and functional diversity of the peri-LC we identified here presents unique opportunities for altering NE function in naturalistic behavior, cognition, motivated behaviors, physiological stress, as well as the many disease-related behavioral states previously associated with the LC. Notably, while peri-LC^GABA^ projections outside of the brainstem are sparse, our study does not explore the possible influence of peri-LC populations on other brainstem areas, which may influence the behaviors we observed.

Our findings presented here identify diverse peri-LC^GABA^ neurons at molecular, cellular, and systems levels, providing avenues for future studies to better understand how these newly identified peri-LC populations function in a host of ethologically relevant behaviors including exploration, feeding, odor and pain processing, decision-making, cognition, and associative learning. We isolated unique peri-LC neuronal groups which may act to integrate ascending and descending LC^NE^ control via both direct fast neurotransmission and diverse arrays of neuropeptide GPCR systems. The expression of several genetically defined subpopulations with differing functional properties represents a subset of putative peptidergic integrators of arousing stimuli and modulators of arousal states mediated via the LC^NE^ system. Further heterogeneity exists in the peri-LC and LC^NE^ populations in terms of transcription factors, ion channels, and other markers, possibly revealing additional layers of interesting LC system complexity. Sub-populations of peri-LC^GABA^ neurons may thereby form distinct modular components of the extended LC^NE^ system, impacting noradrenergic tone, physiological arousal, cognition, and avoidance behavior.

## Supporting information

Supplementary Table S1 - Sexually Dimorphic Genes

Supplementary Table S2 - All Genes of Interest

Supplementary Materials - Pathology, Neurotransmission, and GPCR-related genes

## Methods

### Animals

Adult wild-type C57BL/6J (Jackson Laboratory #000664), vGAT-IRES-Cre (*Slc32a1^tm2(cre)Lowl^*/J; Jackson Laboratory #028862)^86^, pENK-IRES-Cre (*B6;129S-Penk^tm2(cre)Hze^/J*; Jackson Laboratory #025112), Tac1-IRES-Cre (B6;129S-Tac1^tm1.1(cre)Hze^/J; Jackson Laboratory # 021877) and prepronociceptin (*Pnoc*)-IRES-Cre (*Pnoc^tm1.1(cre)Mrbr^*/J; Jackson Laboratory # 034278) ^87^ mice were group housed, given access to food and water *ad libitum*, and maintained on a 12:12 hr reverse light:dark cycle (lights off at 9:00 AM and on at 9:00 PM). All animals were kept in an isolated and sound-attenuated holding facility within the lab and adjacent to behavior rooms one week prior to surgery, post-surgery, and throughout the duration of the behavioral assays to minimize stress. Males and female mice were used in all experiments. All procedures were approved by the Animal Care and Use Committee of Washington University and the University of Washington and conformed to US National Institutes of Health guidelines.

### Stereotaxic Surgeries

After the animals were acclimated to the holding facility for at least seven days, the mice were anesthetized in an induction chamber (2% isoflurane) and placed into a stereotaxic frame (Kopf Instruments, Model 1900) where they were maintained at 1-2% isoflurane. Mice were then injected using a Nanoject (Drummond Scientific #3000204 and #3000207) with a glass tip of 7- 20 μm diameter. Mice were injected in the peri-LC (AP -5.40 mm, ML +0.75 mm, DV -3.70 mm relative to bregma) with three bursts of 9.2 nL each (3 minutes apart; 27.6 nL total) at a rate of 23 nL/s. Injections were unilateral for tracing and fiber photometry experiments; bilateral for optogenetic or chemogenetic experiments. The needle was slowly removed from the brain 10 min after cessation of injection to allow for diffusion. For fiber photometry and optogenetic experiments, mice were also implanted with a 400-µm fiber optic (Doric Inc., MFC_400/430- 0.48_MF2.5_FLT) or custom-made optical fibers of ≥70% optical efficiency in the same surgery. For mice in the peri-LC photostimulation study to examine effect on prefrontal cortex (PFC) and basolateral amygdala (BLA) NE levels, in addition to the peri-LC injections as before, 500nl of the virus expressing the fluorescent NE sensor (GRAB_NE2m) was injected in the BLA (AP - 1.45, ML -3.20 mm, DV -4.75 to -4.50 DV), and 300nL was injected in the PFC (AP +1.80 mm, ML +0.40 mm, DV -2.40 mm), both at 100 nL/min. Fibers were implanted in the peri-LC (AP - 5.40 mm, ML +0.75 mm, DV -3.50 mm), BLA (AP -1.45 mm, ML -3.20 mm, DV -4.50 mm), and PFC (AP +1.80 mm, ML +0.40 mm, DV -2.10 mm), respectively. Mice received carprofen and recovered for at least 6 weeks prior to behavioral testing, permitting optimal expression of the virus. For optogenetic inhibition, fiber optics were placed bilaterally. Fiber optic implants were secured using Metabond (Parkell #S380).

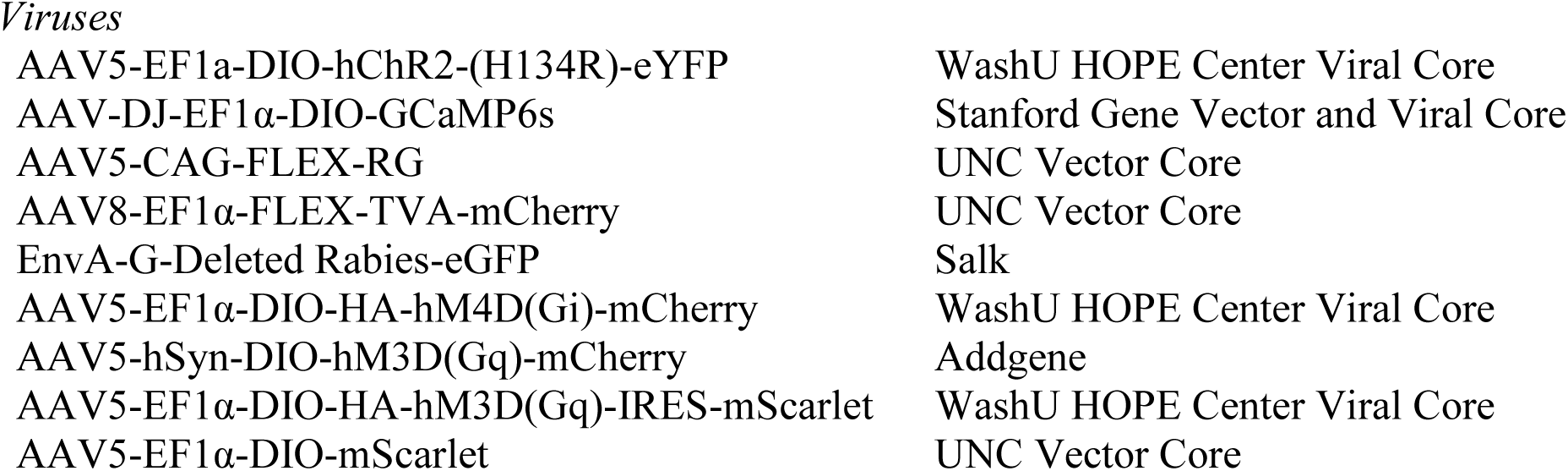
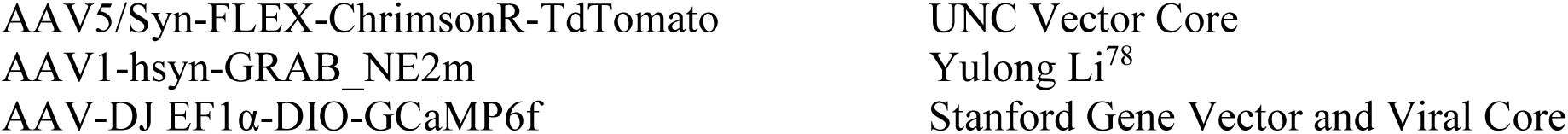

### Immunohistochemistry

Immunohistochemistry was performed as previously described^20^. In brief, mice were anesthetized with sodium pentobarbital and transcardially perfused with 4% paraformaldehyde (PFA), post-fixed overnight in 4% PFA, and cryo-protected in 30% sucrose for at least 24 hours. Brains were then sectioned (30 µm) and placed in 0.1M PB until immunohistochemistry. Free-floating sections were washed in 0.1M PBS for 3 x 10 minutes intervals. Sections were then placed in blocking buffer (0.5% Triton X-100 and 5% natural goat serum in 0.1 M PBS) for 1 hr at room temperature. Sections were incubated with primary antibody (TH – Aves TYH 1:2000; pPDH – Cell Signaling Technology 1:500^80^ at 4°C overnight. After 3 x 10 minute 0.1M PBS, sections were incubated in Alexa Fluor secondary antibodies (1:1000) and NeuroTrace (1:400, 435/455 blue fluorescent Nissl stain, Invitrogen #N21479) for 1 hour, followed by 3 x 10 minute 0.1M PBS then 3 x 10 minute 0.1M PB washes. Sections were then mounted and coverslipped with Vectashield Vibrance HardSet mounting medium (Vector Laboratories, CAT #H-1700) and imaged on Olympus Fluoview 3000 confocal microscope. Animals that did not show targeted expression were excluded. For retrograde rabies tracing, areas were registered using the Paxinos and Franklin brain atlas^88^. For pPDH staining, DREADD+ mice were injected with 1 mg/kg CNO 60 minutes prior to perfusion and cell counts were quantified using QuPath^89^.

### Fluorescent *in situ* Hybridization (FISH)

Animals were anesthetized and rapidly decapitated. Brains were quickly removed and fresh frozen in dry ice, then stored at -80°C. Sections were cut at 20µm, mounted on slides, and stored at -80°C. Sections were fixed in 4% PFA for 15 min, dehydrated in serial ethanol concentrations (50%, 70%, and 100%) and processed with RNAscope (Advanced Cell Diagnostics, cat. No. 320293). Sections were hybridized with the probes listed below. Sections were then counterstained with DAPI and coverslips were applied. Confocal images were obtained on an Olympus FV3000RS microscope; counts were specific to the peri-LC region.

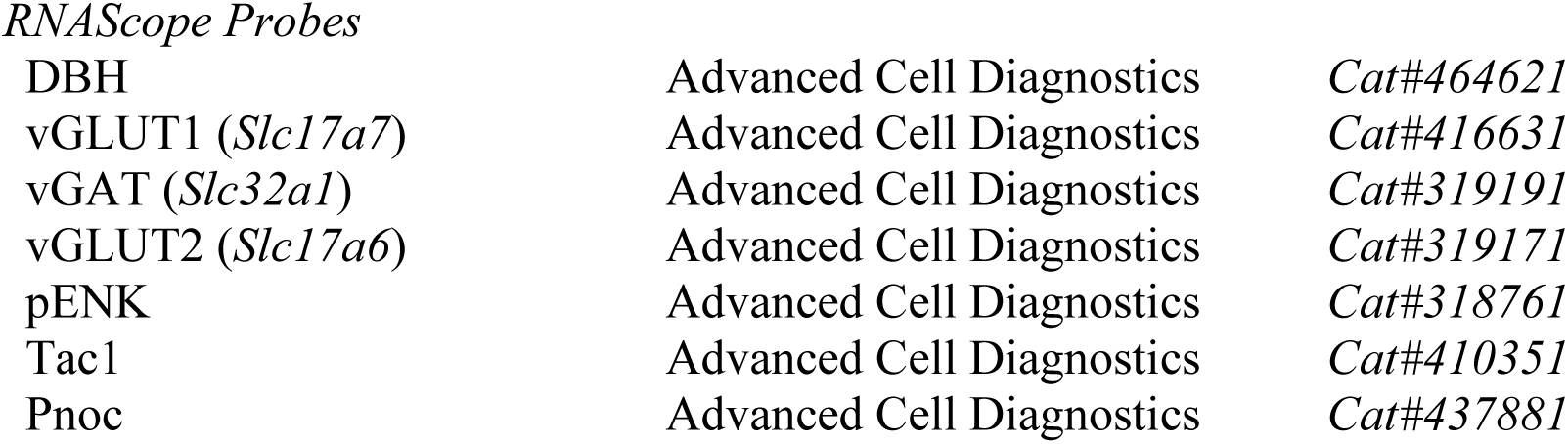

### Slice electrophysiology

For electrophysiology experiments, slices of the LC were prepared from injected mice at least 21 days after stereotaxic surgery. Anesthetized mice underwent cardiac perfusion with ice cold cutting solution (in mM: 75 NaCl, 2.5 KCl, 6 MgCl_2_, 0.1 CaCl_2_, 1.2 NaH_2_PO_4_, 25 NaHCO_3_, 2.5 D-glucose, 50 sucrose; bubbled with 95% O_2_ and 5% CO_2_). The brain was then submerged in cutting solution and placed on a VT1200 vibratome (Leica), where 240 μm horizontal LC slices were collected. Slices were transferred to a 32°C holding chamber containing oxygenated artificial cerebrospinal fluid (ACSF; in mM: 126 NaCl, 2.5 KCl, 1.2 MgCl_2_, 2.5 CaCl_2_, 1.2 NaH_2_PO_4_, 21.4 NaHCO_3_, and 11.1 D-glucose; bubbled with 95% O_2_ and 5% CO_2_), where they recovered for at least 1 hour.

Whole cell and cell attached recordings were made using an Axopatch 200B Amplifier (Molecular Devices). Data were acquired with Axograph X (Axograph Scientific) at 5 kHz and filtered to 2 kHz. Cells were visualized with a BX51WI microscope (Olympus). LC^NE^ cells were identified by morphology, location (between the edge of the 4th ventricle and motor neurons of the mesencephalic tract), and the presence of an electrically evoked current (α2-mediated IPSC)^22^. Channelrhodopsin2 expressing peri-LC cells were identified by the presence of a steady state current during a 150 ms light pulse and were medial to LC^NE^ cells. In the recording chamber, slices were continuously perfused at 1.5 - 2 mL per minute with ACSF warmed to 30 +/- 2°C. Pipettes (1.5 - 2.0 MΩ) were pulled from borosilicate glass (World Precision Instruments). For whole cell voltage clamp recordings, cells were held at -60 mV and the internal pipette contained: 135 mM KCl, 0.1 mM CaCl_2_, 2 mM MgCl_2_, 10 mM HEPES, 0.038 mM EGTA, 1 mg/mL ATP, 0.1 mg/mL GTP, 1.5 mg/mL phosphocreatine (pH 7.4, 275 mOsm). Currents were evoked with wide-field stimulation using a 1 ms pulse from an LED (470 nm, ∼0.5-1.0 mW/mm^2^) through a 40x water immersion objective. Paired pulse ratios were measured using a 50 ms interpulse interval. Glutamatergic and GABAergic synaptic components were blocked using DNQX and picrotoxin, respectively. TTX and 4-aminopyridine were used to isolate monosynaptic ChR2-evoked currents. Current clamp recordings were collected (using the same internal solution) to measure action potentials and resting membrane potential of peri-LC cells. For cell attached recordings, the intracellular pipette solution contained: 126 mM NaCl, 2.5 mM KCl, 1.2 mM MgCl_2_, 2.5 mM CaCl_2_, and 1.2 mM NaH_2_PO_4_. Firing rate was measured in 500 ms bins, with two cells that did not recover firing excluded from analysis.

### Brain clearing and imaging LC and peri-LC

VGAT-Cre mice were injected with 300 nl of AAV5-EF1α-DIO-ChR2-(H134R)-eYFP (1x10^13^ vg/mL) at -5.40 AP, +0.75 ML, -4.0 to -3.7 DV from bregma with a lateral-facing beveled syringe. After 5 weeks, mice were euthanized, perfused with phosphate buffered saline and paraformaldehyde, and the brains were collected. Tissue clearing was modified from iDISCO+ protocol^90^. Briefly, brains were dehydrated with methanol, delipidated with dichloromethane, and bleached with hydrogen peroxide solution. After rehydration and blocking, brains were incubated with 1:1000 chicken anti-GFP antibody (Aves GFP-1020) and 1:1000 mouse anti-tyrosine hydroxylase antibody (BioLegend 818001), with boosters over 5 days for a final concentration of 1:200. After washing, secondary antibodies were added: 1:500 goat anti-chicken IgG Alexa Fluor 647 (Invitrogen A21449) and 1:500 goat anti-mouse Alexa Fluor 488 (Life Technologies A11001), with boosters over 5 days for a final concentration of 1:100. After washing and dehydration, brain clearing and refraction index matching were performed with dibenzyl ether. Brains were imaged with a light sheet microscope (UltraMicroscope II, LaVision Biotec), and imaging data were analyzed using ImageJ and Arivis Vision4D (3.0.0).

### Single-nucleus RNA sequencing

Adult mice were euthanized with 0.01 mL/kg of phenytoin/pentobarbitol solution, and perfused transcardially with cold NMDG-aCSF (96 mM NMDG, 2.5 mM KCl, 1.35 mM NaH_2_PO_4_, 30 mM NaHCO_3_, 20 mM HEPES, 25 mM glucose, 2 mM thiourea, 5 mM Na+ ascorbate, 3 mM Na+ pyruvate, 2 mM N-acetyl-cysteine, 0.5 mM CaCl_2_, 10 mM MgSO_4_, 0.6 mM glutathione-ethyl-ester; pH 7.35-7.40, 300-305 mOsm, oxygenated with 95% O_2_ and 5% CO_2_) with the following inhibitors: 500 nM TTX, 10 μM APV, 10 μM DNQX, 5 μM actinomycin, 22.7 μM anisomycin. Extracted brains were sliced using a vibratome (Leica, VT1200), the locus coeruleus and its neighboring regions dissected out, and the tissue flash frozen on dry ice and stored at -80°C. Tissue from 5-6 mice was pooled for each sample; male and female samples were processed separately. Later, coerulear sections were lysed, homogenized with glass pestle, filtered via a 40 μm strainer, centrifuged initially to remove cellular debris, and again in differing density solvents (50% and 30% OptiPrep) in which the pelleted nuclei were collected. Nuclei were resuspended in PBS as needed for >400 nuclei/μL.

The cDNA libraries were constructed using the Chromium Next GEM Single Cell 3’ Reagents Kits v3.1 Dual Index, 10x Genomics. Briefly, ∼10,000 nuclei were load into the chip, single nuclear mRNAs were captured by barcoded beads, and the transcripts were reverse transcribed. The resulting cDNAs were then PCR amplified, fragmented, ligated with adaptors, and added with sample index. To minimize batch effects, experimenters were well-practiced, timing was kept the same, and all reagent lots were the same. cDNA libraries were sequenced on Illumina Hiseq 2x150bp and NovaSeq PE150, and sequence alignment to mouse genome done with 10x Genomics Cell Ranger Count v6.1.2 pipeline.

Integration of data sets, clustering, and differential gene expression analysis were performed using Seurat v4.0^91^ and R. Nuclei of potentially low quality (total UMI<700 or total UMI>15,000, or percentage of mitochondrial genes >0.1%) were removed from downstream analysis. Suspected doublet cells were computationally removed using DoubletDecon 1.02^92^. Cells were clustered by weighted-nearest neighbor analysis of normalized gene expression and these clusters were validated using known markers for cell types in the brain^91^. Dimensions and resolution were determined by elbow plots and ChooseR^63^. Differentially-expressed genes between clusters were identified using a non-parametric Wilcoxon rank sum test^93^. Gene lists were acquired and modified from ontology lists at the Mouse Genome Database (MGD), Jackson Laboratory, Bar Harbor, Maine (informatics.jax.org) (December 2021)^94^.

### PIXEL-seq

Pixel-seq was performed as previously described in a recent report^69^. Briefly, polyacrylamide gels of <5mm thickness were cast and crosslinked on a 40 mm round coverslip (#1.5; Warner Instruments 64–1696) in an anaerobic chamber. The gels were assembled into a FCS2 flowcell (Bioptechs) and 5′ phosphorothioate-modified primers were covalently grafted onto the gel. Next, 368-bp DNA templates (Integrated DNA Technologies)were hybridized onto the gel by washing on 8-12 pm mix (0.6-0.8 million features/mm^2^) and incubated at 75°C for 2 min, cooling to 40°C, and washing with 500 μL amplification buffer. Each template contained 24-bp random spatial indices, which serve as spatial barcodes, a 20-bp poly(T) probe, which captures RNAs, and two Taq1 restriction sites, which are digested to expose the poly(T) probes for RNA capture. Following this, a 150 μL Taq polymerase mixture was used to synthesize and anchor the first-strand DNAs to the gel by incubating at 74°C for 5 min.

Polonies were amplified to reduce gaps between the polony templates, then digested with TaqI, and polony sequencing using sequencing-by-synthesis (HiSeq SBS kit v4; Illumina FC-401- 4002) was used to build a spatial index of barcode coordinates. For more details see ^69^.

A 10μm-thick section of fresh-frozen brain containing the LC was dissected using a cryostat (Thermo Scientific NX70) and placed on the gel in a flat layer. The tissue sections were then gently immersed in 50 μL hybridization buffer (6x SSC (Invitrogen AM9763), 2U/μL RNaseOUT (Thermo Fisher 10777019) and incubated at 23°C for 15 minutes, hybridizing them to the capture probes adhered to the gel. To create cDNAs,*in situ* reverse transcription was performed in a reaction chamber by incubating for 42°C for 1 hr in 100 μL reverse transcription (RT) mixture (5 μL Maxima H- reverse transcriptase (Thermo Fisher, EP0753), 20 μL 5x Maxima RT buffer, 20 μL 20% Ficoll PM-400 (Sigma-Aldrich, F4375), 10 μL 10 mM each dNTPs, 5 μL 50 μM template-switching oligo (Qiagen, 339414YCO0076714), 2.5 μL RNAseOUT (40 U/μL), and 37.5 μL H2O). After removing the reaction buffer, tissues were digested using proteinase K (10 μL proteinase K (Qiagen, 19131) in 90 μL PKD buffer (Qiagen, 1034963)) at 55 °C for 30 min and then washed away. To recover the spatially barcoded cDNAs, the chambers were incubated for 15 min at 65°C with 70 μL second-strand mix (7 μL 10 × isothermal amplification buffer (NEB, B0537), 7 μL 10 mM each dNTP mix, 3.5 μL 10 μM TSO primer, 0.5 μL 20 mg/mL BSA (NEB, B9000), 3 μL BST2.0 WarmStart DNA Polymerase (NEB, M0538), and 49 μL H2O), then the cDNAs eluted into a tube. To introduce UMIs, ∼35 μL sample was mixed with 65 μL UMI incorporation mix (50 μL 2x Q5 Ultra II master mix (NEB, M0544), 2.5 μL 10 μM UMI primer, and 12.5 μL H2O), then denatured at 95 °C for 30 s, annealed at 65 °C for 30 s, and extended at 72°C for 5 min. 12 cycles of PCR amplification of the cDNA library were performed as previously described^69^. 1ng DNA was then used for sequencing library construction using a Nextera XT kit (Illumina FC-131-1024) following the manufacturer’s protocol. cDNA libraries were sequenced on NovaSeq PE150.

Transcripts were mapped to the spatial barcode map by processing the FASTQ files. After using Flexbar^95^ to extract spatial barcode and UMI sequences, Bowtie^96^ was used to map index sequences to the spatial barcode map, allowing up to 2 mismatches. STARv2.7.0^97^ was used to align paired-end reads of mapped indices to a reference mouse transcriptome (GRCm38). Sequencing reads with the same transcriptome mapping locus, UMI, and spatial barcode were collapsed to unique records for subsequent analysis. The cell segmentation was performed as previously described^69^.

We next performed spatial pattern genes analysis using SpatialPCA^98^. ∼1200 genes were identified. For the female and male LC clustering, SCTransform normalization was applied in Seurat using 2,000 highly variable genes. We choose to combine the spatial pattern genes and highly variable genes as features for the subsequent analysis. After Principal Component Analysis (PCA), 20 PCs were used in the UMAP visualization, and the clustering graph was generated with a resolution of 0.6. Marker genes for each cluster were identified by Seurat using a Wilcoxon test and then compared with the consensus in a published mouse brain cell type atlas^99^.

Segmented LC cells were annotated with a snRNA-seq dataset (GSE121891)^100^ as a reference to predict cell type compositions using scvi-tools^101^. The top 3,000 variable genes were selected for the model training. Raw gene counts for training and testing datasets were scaled to 10^4^. The training and prediction were run at max_epochs = 500. Cell types of segmented cells were determined by the predicted cell types with the highest ratios. For the outcome of Fig.4b, the threshold of trusted cell type was set as 0.5.

### Fiber photometry

Fiber photometry recordings were performed as previously described^87^. Briefly, an optic fiber was attached to the implanted fiber by a ferrule sleeve, then GCaMP6s was stimulated by two LEDs, a 531-Hz sinusoidal light (Thorlabs M470F3), bandpass filtered at 470 ± 20nm, and a 211-Hz sinusoidal light (Thorlabs M405FP1), bandpass filtered at 405 ± 10nm. (Filter cube: Doric FMC4; LED driver: DC4104). The 470 nm signal evokes Ca^2+^-dependent emission, while the 405 nm signal evokes Ca^2+^-independent isosbestic control emission. Prior to recording, a 180s period of GCaMP6s excitation with both light channels was used to remove the majority of baseline drift. Laser intensity at the optic fiber tip was adjusted to ∼50 μW before each day of recording. GCaMP6s fluorescent signal was isolated by bandpass filtering (525 ± 25nm), transduced by a femtowatt silicon photoreceiver (Newport 2151), and recorded by a real-time processor (TDT RZ5P). The envelopes of 531 Hz and 211 Hz signals were extracted in real time by the TDT program Synapse at a sampling rate of 1017.25 Hz. The 565nm LED was used for peri-LC photostimulation,with light intensity adjusted to ∼2.5µW at the optical fiber tip, while GRAB_NE signals were simultaneously recorded from other cables from the PFC and BLA.

Fiber recordings were analyzed using custom MATLAB scripts available on Github. The isosbestic signal was subtracted from the calcium-dependent signal, then baseline drift due to slow photobleaching artifacts was corrected by fitting a double exponential curve to the raw trace. The baseline-corrected signal was low-pass filtered using a Butterworth filter with cutoff set at 10 Hz. The filtered signal was then normalized by subtracting the median signal value and dividing by the absolute median value of the isosbestic signal. The correct and normalized photometry trace was z-scored relative to the mean and standard deviation of the signal over the entire trial. The mean z-score during comparable periods of time determined by the stimulus duration (when applicable) or the experimentally-determined duration of a neural response (see Supplementary Table S1) were compared using paired t-tests. For determination of neural activity during movement, mice were tracked using Ethovision 10; movement was defined as all samples with movement ≥10cm/s (in open field) or ≥5cm/s (in zero maze) for ≥0.5s. The average activity during movement and non-movement per session were then compared using paired t-tests.

### Two-photon slice calcium imaging

Brains were obtained from Dbh-Cre mice >4 weeks after injection of AAV-DJ-EF1α-DIO- GCaMP6f in the LC, and freshly sliced at 250 μm in NMDG solution (92mM NMDG, 20mM HEPES, 25mM D-glucose, 30mM NaHCO_3_, 1.45mM NaH_2_PO_4_, 2.5mM KCl, 5mM Na ascorbate, 3mM Na pyruvate, 2mM thiourea, 0.5mM CaCl_2_, 2M MgSO_4_) using vibratome. Brain slices were placed on imaging slide and secured with a grating anchor. Artificial cerebrospinal fluid solution (ACSF, 126mM NaCl, 2.5 KCl, 1.2mM NaH_2_PO_4_, 1.2 mM MgCl_2_, 11mM D-glucose, 18mM NaHCO_3_, 2.4mM CaCl_2_) is delivered to the slide by arotatory pump (Gilson Minipuls) and through an in-line warmer (Warner Instrument TC-324C) to maintain at 31-34°C. Three-minute treatments with ACSF, DAMGO (Tocris 1171), Leu-enk (Thermo scientific J61408.MB), and nociceptin (Tocris 0910) were counterbalanced and washed out for 3 min before the subsequent treatment. KCl treatment was for 1 min at the end of experiment. Suite2p was used to detectROIs, and Excel for further analysis. The median signal from 1.5 min to 2.5 min after drug perfusion was used for comparison to the 1 min baseline prior to perfusion.

### Behavior

#### Odor delivery

Odor delivery was performed in a polyethylene chamber approximately 12 x 12 x 24 cm, out of which air was continuously vacuumed at a rate of 2L/min. To minimize odor release into the room, the chamber was placed in a fume hood and the vacuumed air was passed through a carbon filter system. Mice were habituated to the room and chamber three times before odor was delivered and were habituated to the room for 30 min before testing. The chamber was wiped with 70% ethanol between trials, which fully evaporated before the next animal was tested. Odor was delivered for 30s periods at a rate of 0.5 L/min. Tubing was changed between odorants to minimize cross-contamination. Bobcat urine (predatorpeestore.com) was used undiluted. Opposite sex urine was collected from unstressed mice an hour before experiments began. 2-PE (Sigma 77861) was diluted 1:10 in mineral oil. Peppermint oil (Sigma Aldrich 77411) was diluted 1:100 in mineral oil. 2MT (TCI M0285) was diluted 1:50 in mineral oil.

#### Food consumption

Feeding assays were performed in a square plexiglass arena (27cm × 27cm). Mice were habituated to the room and chamber at least 3 times before testing and the room 30 min before trials. For food-deprived conditions, mice were food deprived for 24 h before trial. For highly-palatable food, mice were habituated to the food item one week before trial to eliminate food neophobia. Food items were placed in the corner of the arena and mice were introduced to the area for 30 min. To record the amount of food consumed, food was measured before and after the assay; the bottom of the chamber was scraped after each trial to collect all crumbs resulting from chewing without consumption. When multiple stimulation or inhibition conditions were tested, they were performed in a counterbalanced fashion across mice. For DREADD experiments, clozapine N-oxide (CNO) (1 mg/kg, Enzo Life Sciences, Cat#BML-NS105) was administered to both control and DREADD+ animals 30 minutes prior to their individual test time.

#### Threat Conditioning

Pavlovian threat conditioning was performed in Med Associates Fear Conditioning Chambers (NIR-022MD). This equipment consisted of a conductive grid floor inside a 29.53cm L x 23.5cm W x 20.96cm H chamber inside of a soundproof box lit by an infrared light. Stimuli occurred at random timing, at least 110-140s apart. On day 1, mice were exposed to six 20s tones without an associated shock. On day 2, mice were conditioned to six 20s tones co-terminating with 1-s 0.5mA shocks. On Day 3, mice were presented with six 20s tones in the absence of shock.

#### False food

False food testing was performed as previously described^102^. Mice were food deprived for 24 hours before testing, and were habituated to the room for 30 minutes before testing. Mice were then placed in a dimly-lit (25 lux) plexiglass arena (27 cm x 27 cm) with a foam plug shaped like a piece of chow placed in the middle. Videos were scored for attempts to eat the false food.

#### Novel object

Novel object testing was performed as described previously^20^. Briefly, mice were habituated to the room for 30 minutes before testing. Mice were then placed in a dimly-lit (25 lux) plexiglass arena (27 cm x 27 cm) with two identical objects (test tubes or petri dishes; counterbalanced) fixed at the corners. Mice were given ten minutes to explore the objects, then returned to the homecage for one hour, then were placed in the arena with one familiar object and one novel object. Objects were thoroughly cleaned with ethanol and allowed to dry between trials. Videos were scored for interactions with each object.

#### Elevated Zero Maze (EZM)

EZM testing was performed as previously described^20^. The EZM (Harvard Apparatus) was made of grey plastic, 200 cm in circumference, comprised of four 50 cm sections (two opened and two closed). The maze was elevated 50 cm above the floor and had a path width of 4 cm with a 0.5 cm lip on each open section. Mice were connected to fiber optic cables, positioned head-first into a closed arm, and allowed to roam freely for 20 min. Open arm time was the primary measure of anxiety-like behavior. For optogenetic experiments, animals received 4 Hz (5 ms pulse width) constant photostimulation (2mW light power). For DREADD experiments, 1 mg/kg CNO was administered to both control and DREADD+ animals 30 minutes prior to their individual test time.

#### Open Field Test (OFT)

OFT was performed as described previously^20^ in a plexiglass arena (50 cm x 50 cm). Center zone was defined as the middle 50% of the arena size. Mice were connected to fiber optic cables, positioned head first into a closed arm, and allowed to roam freely for 20 min. For optogenetic experiments, animals received 4 Hz (5 ms pulse width) constant photostimulation (2 mW light power). For DREADD experiments, 1 mg/kg CNO was administered to both control and DREADD+ animals 30 minutes prior to their individual test time. Center time was the primary measure of anxiety-like behavior.

#### Real-Time Place Preference (RTPP)

Mice were placed into a custom-made unbiased, balanced two-compartment conditioning apparatus (52.5cm x 25.5cm x 25.5cm) as previously described^20^ and allowed to freely roam the entire apparatus for 20 min. Entry into one compartment triggered constant photostimulation (0- 20 Hz; 2 mW light power) while the animal remained in the light-paired chamber. Entry into the other chamber ended the photostimulation. The side paired with photostimulation was counterbalanced across mice and between days. Time spent in each chamber and total distance traveled was measured using Ethovision 10 (Noldus).

#### Light-dark box

Light-dark box was performed in a RTPP chamber where one side was brightly lit (1000 lux) and the other side was dimly lit (5 lux). Animals were allowed to freely move throughout the chamber for 20 minutes. Time spent in the light side was the primary measure of anxiety-like behavior. For optogenetic experiments, animals received 4 Hz (5 ms pulse width) constant photostimulation (2mW light power). For DREADD experiments, CNO was administered to both control and DREADD+ animals 30 minutes prior to their individual test time.

#### Foot pinch

Animals were lightly anesthetized at 1.5% isoflurane and placed in a stereotax. Their pupil size was monitored under infrared lighting^41^. Mice were pinched on their contralateral paw for 10s.

#### Tail flick

The tail-flick assay was performed as previously described^103^. 2-4 cm of the mouse’s tail was immersed into 54°C water and the time between immersion and withdrawal was recorded, up to a maximum of 10s in order to prevent tissue damage. For DREADD experiments, DREADD+ and control mice were injected with 1mg/kg CNO after the initial test (time 0) and tested every 10 min for one hour, then again at 90 min.

#### Hot plate

The hot-plate assay was performed as previously described^104^. Mice were placed on a 55°C hot plate and the time for a nocifensive response (paw withdrawal or licking, stamping, leaning, or jumping) was recorded, up to a maximum of 30s in order to prevent tissue damage. All mice were injected with 1mg/kg CNO 30 min before their individual test time.

### Statistical analyses

All summary data are expressed as mean ± SEM. Statistical significance was taken as *p < 0.05, **p < 0.01, ***p < 0.001, ****p < 0.0001, as determined by two-tailed Student’s t-test (paired and unpaired), two-way repeated measure analysis of variance (ANOVA) followed by Tukey’s multiple comparison test, two-tailed Wilcoxon matched-pairs signed rank test, and Friedman repeated measures ANOVA followed by Dunn’s multiple comparison test. Statistical analyses were performed in GraphPad Prism 9. For sequencing experiments, analyses were performed as described in the corresponding methods section. Summary data is available in Supplementary Table S1.

### Data availability

All data are available in the manuscript or the supplementary materials. snRNAseq data is deposited in in the NCBI Gene Expression Omnibus (GEO) database with accession number GSE201998; sample GSM6086497 – LC_FEMALE_01 is included for the sake of completeness but it is poor quality and we do not recommend using it. PIXEL-seq data is deposited in GEO database with accession numbers GSE244659. The full behavioral dataset supporting the current study are available from the corresponding author upon reasonable request.

### Code availability

Custom MATLAB analysis code was created to appropriately organize, process, and combine photometry recording data with associated behavioral data. Custom R scripts were written to analyze single nucleus RNA sequencing datasets. Code supporting the current study are available from the corresponding author upon reasonable request.

## Acknowledgments

We thank Dylan Blumenthal, Michelle Chung, Megan Votoupal, Taylor Hobbs, Carina Pizzano, and Valerie Lau for animal colony maintenance; Christine Stander and Azra Suko for laboratory management and support; the Bruchas laboratory and UW NAPE Center for discussions; Marcus Basiri and the Stuber Lab for assistance with single-cell sequencing; Jane Chen and Richard Palmiter for mouse lines and assistance with single-cell sequencing. We thank Chris Stander, K. Deisseroth, the Washington University HOPE Vector Core, and the University of North Carolina (UNC) Vector Core for viral constructs, prep and packaging.

## Funding

This study was supported by the National Institute of Mental Health (ATL, F31- MH122033; RCS, F31-MH117931; LS, R41-MH130299; CPF & KB, R21-MH123085; MRB, R01-MH112355), the National Institute on Drug Abuse (LL, K99-DA054446; SAG, R00- DA045662, UG3-DA053802, & R01-DA054317; GDS, R37-DA032750 & R01-DA038168; CPF, R01-DA035821; LG & MRB, R61- DA051489; MRB, P30-DA048736), the National Cancer Institute (LG, UG3-CA268096), the National Institute of General Medical Sciences (ATL, T32- GM008151), the Foundation for Anesthesia Education and Research (LL), and the NARSAD Young Investigator Award 27082 (SAG). The content is solely the responsibility of the authors and does not necessarily represent the views of the National Institutes of Health nor any other funding institution.

## Author contributions

ATL, LL, and MRB conceptualized and designed the entire study. ATL, LL, and MMM performed stereotaxic surgeries. ATL, MMM, TB, BAW, STT, EML, and ANR performed behavioral experiments. ATL, LL, MMM, TB, BAW, STT, EML, and ANR performed histology. ATL, LL, XF, LS, and LG performed Pixel-seq experiments. KB and KSG performed electrophysiological experiments. ATL, LL, and TB performed fiber photometry experiments. ATL, LL, and RCS performed single-nucleus RNA sequencing experiments and analysis. ATL, LL, MMM, and KB performed statistical analysis. LL and ADM performed tissue clearing and imaging. ATL, LL, XF, MMM, KB, and MRB made the figures. ATL, LL, and MRB wrote the manuscript with collective input from all authors. SAG supervised tissue clearing and imaging. GS supervised single-nucleus sequencing. CPF supervised electrophysiological experiments. LG supervised Pixel-seq experiments. MRB oversaw, provided resources and supervised the entire study.

## Competing interests

Two patents on polony gel stamping and Pixel-seq have been filed by the University of Washington and TopoGene Inc. L.G. and L.S. are co-founders of TopoGene, which manufactures polony gels used in our approaches herein.

## Supplementary Information

**Supplementary Table S1. – Statistical Summary.xlsx**

Summary of statistical tests, photometry time course, rabies tracing results, and scRNAseq experimental details.

**Supplementary Table S2.** – All Genes of Interest.xlsx

Relative expression of genes in clusters of cells in all neurons, NE neurons, and GABA neurons.

**Supplementary Table S3.** – Sexually Dimorphic Genes.xlsx

Sexually dimorphic genes in all neurons, NE neurons, and GABA neurons are shown in order of decreasing significance. Sign of fold-change is positive when higher in male samples.

**Supplementary Video 1**

The video shows a representative 3D rendering in a cleared brain tissue of anti-tyrosine hydroxylase-stained LC neurons (green) and the neighboring anti-EYFP stained peri-LC GABAergic neurons (red) expressing ChR2-eYFP in a Cre-dependent manner in a vGAT-Cre mouse.

---

**Extended Data Fig. 1.**
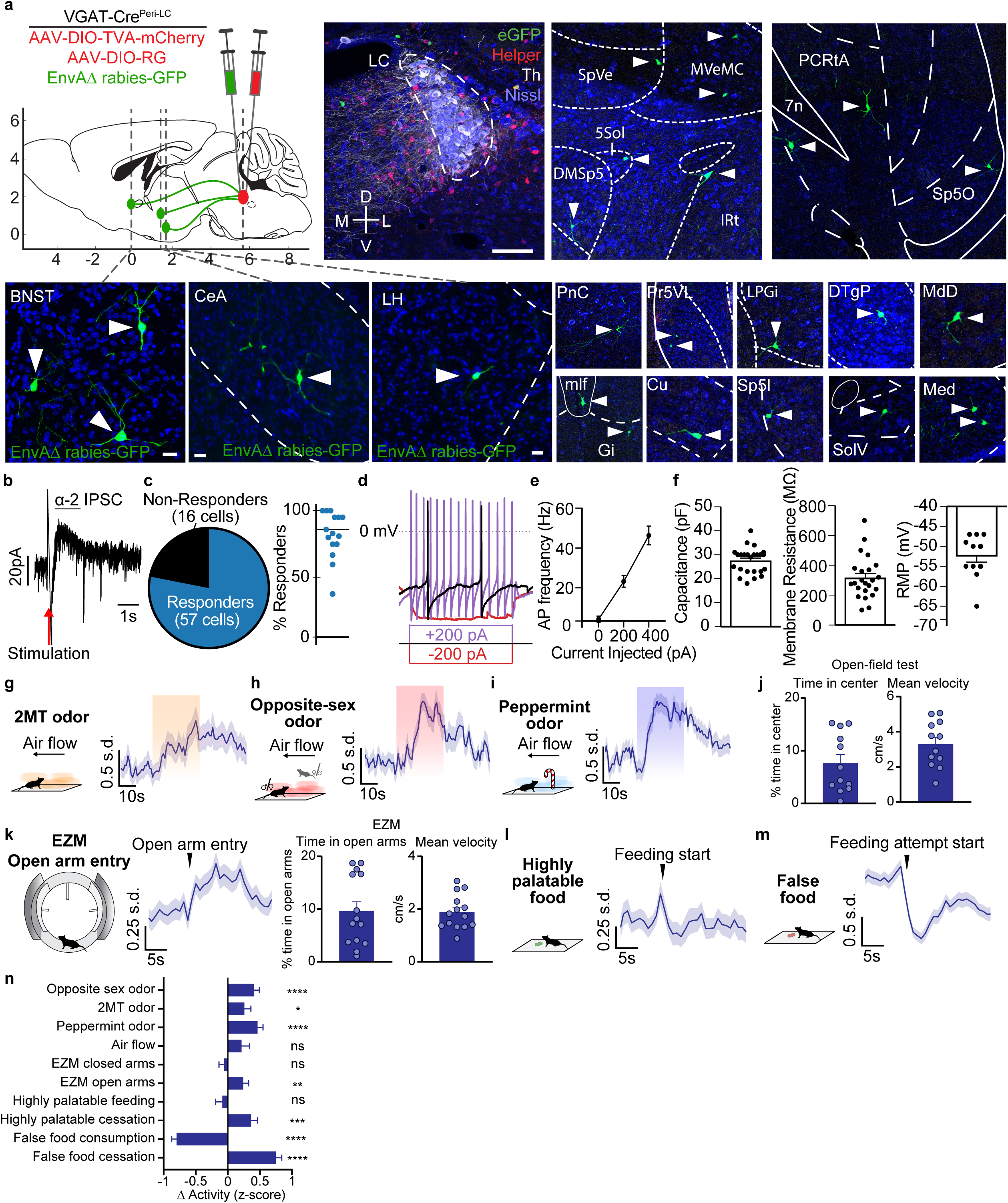
*In vitro* and *in vivo* identification and characterization of pericoerulear GABAergic neurons. **(a)** Viral schematic for rabies tracing of monosynaptic inputs to peri-LC^GABA^ neurons. Starter cells in peri-LC express TVA-mCherry and rabies-GFP (scale bar: 100 μm in LC; 20 μm in input regions). **(b)** Electrically evoked α2-mediated outward current recorded from LC^NE^ neuron. **(c)** 78% of LC^NE^ neurons displayed measurable fast IPSCs, though the success rate appeared dependent on injection specificity as 6 of 14 injection sites had 100% response rate (each data point represents one hemisphere, line shown is median; injections that were completely off target and/or had 0% response rate were excluded). **(d)** Properties of peri-LC GABA neurons. ChR2-expressing peri-LC neurons were identified by the presence of eYFP expression and a steady state response to a 150 ms light pulse (not shown). Representative trace of a current clamp recording with steps of 200 pA current injection. **(e)** Peri-LC^GABA^ neuron firing rate upon current injection (mean values ± SEM). **(f)** Peri-LC^GABA^ neuron capacitance, membrane resistance, and resting membrane potential (RMP) (mean values ± SEM). **(g)** Peri-LC^GABA^ neurons increase activity during 30s exposure to 2MT. **(h)** Peri-LC^GABA^ neurons increase activity during 30s exposure to odor of opposite-sex mice. **(i)** Peri-LC^GABA^ neurons increase activity during 30s exposure to peppermint oil odor. **(j)** Summary of exploration in open-field test. **(k)** Peri-LC^GABA^ neuronal activity increases upon open arm entry in an elevated zero maze. **(l)** Peri-LC^GABA^ neurons are not affected by the consumption of highly-palatable food. **(m)** Peri-LC^GABA^ neurons decrease activity during attempts to eat a false food pellet. **(n)** Summary of fiber photometry experiments. Bars represent change in activity before vs. after events. Periods of comparison were 30s for odor experiments and air flow, 5s for maze exploration, and 10s for feeding. n=12 animals for photometry experiments. \**p* < 0.05, ** p<0.01, *** p<0.001, ****p<0.0001 by paired two-tailed *t*-test. Error bars indicate SEM.

**Extended Data Fig. 2.**
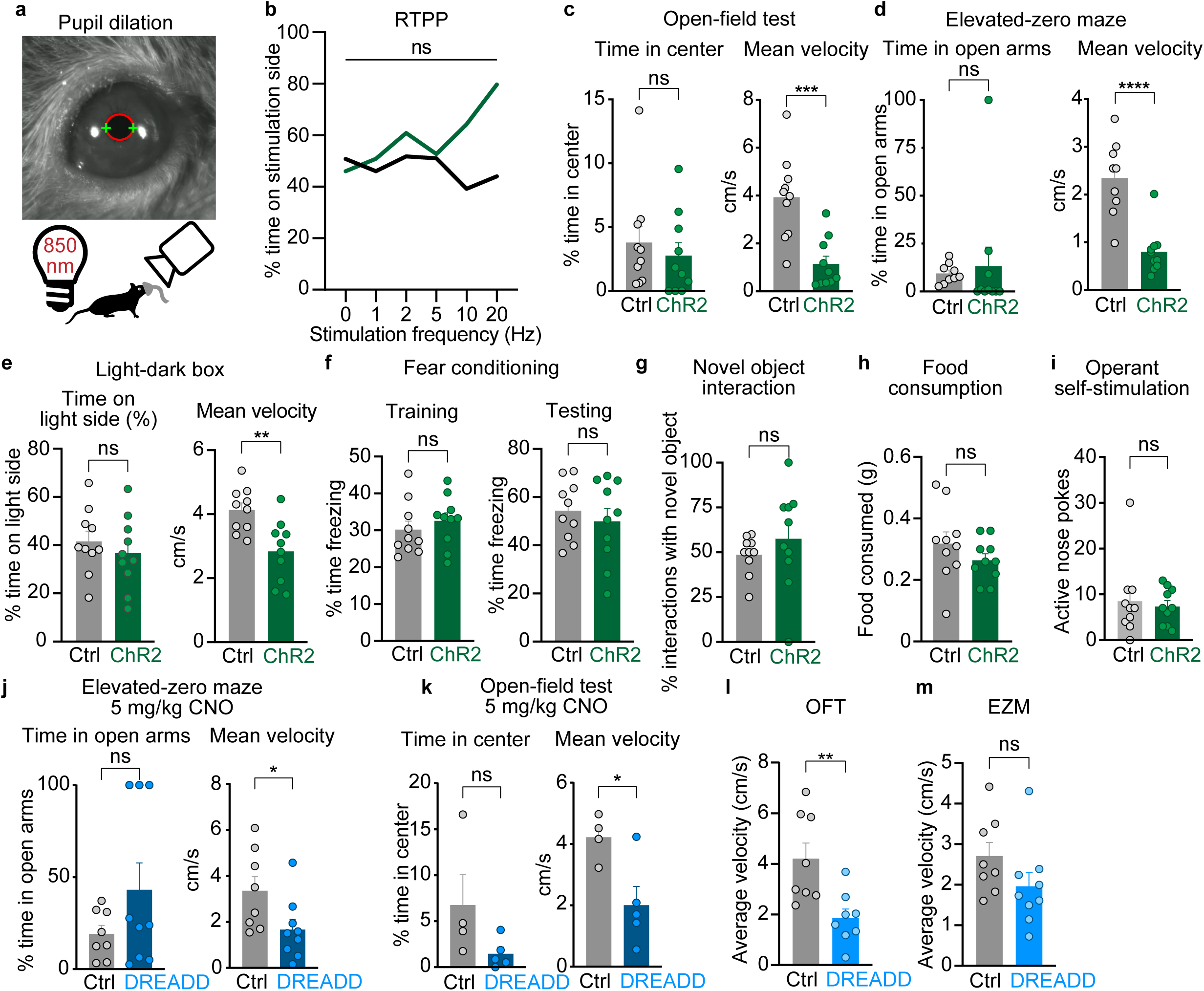
Selective manipulation of pericoerulear GABA neurons modulates physiological arousal and exploratory behaviors. **(a)** Schematic and image of pupil dilation experiment. **(b)** Photostimulation did not cause a real-time place preference. **(c-e)** Photostimulation (2 mW, 4 Hz, 5 ms pulse width) affected movement, but not anxiety-like behavior, in open-field test (c), elevated-zero maze (d), and light-dark box (e). **(f)** Photostimulation (2 mW, 4 Hz, 5 ms pulse width during 20s presentation of cue during fear conditioning) did not affect acquisition of fear response. **(g)** Photostimulation (2 mW, 4 Hz, 5 ms pulse width) did not affect novel object interaction. **(h)** Photostimulation (2 mW, 4 Hz, 5 ms pulse width) did not affect food consumption in food-deprived state. **(i)** Animals did not preferentially nose-poke to receive photostimulation (2 mW, 4 Hz, 5 ms pulse width). **(j, k)** Chemogenetic inhibition (5 mg/kg CNO) of peri-LC^GABA^ neurons caused behavioral arrest in elevated-zero maze and open-field test. **(l)** Chemogenetic inhibition (1 mg/kg CNO) of peri-LC^GABA^ neurons decreased movement in open-field test. **(m)** Chemogenetic inhibition (1 mg/kg CNO) of peri-LC^GABA^ neurons did not affect movement in elevated-zero maze. \**p* < 0.05, ** p<0.01, ***p<.001, ****p<0.0001 by unpaired two-tailed *t*-test. Error bars indicate SEM.

**Extended Data Fig. 3.**
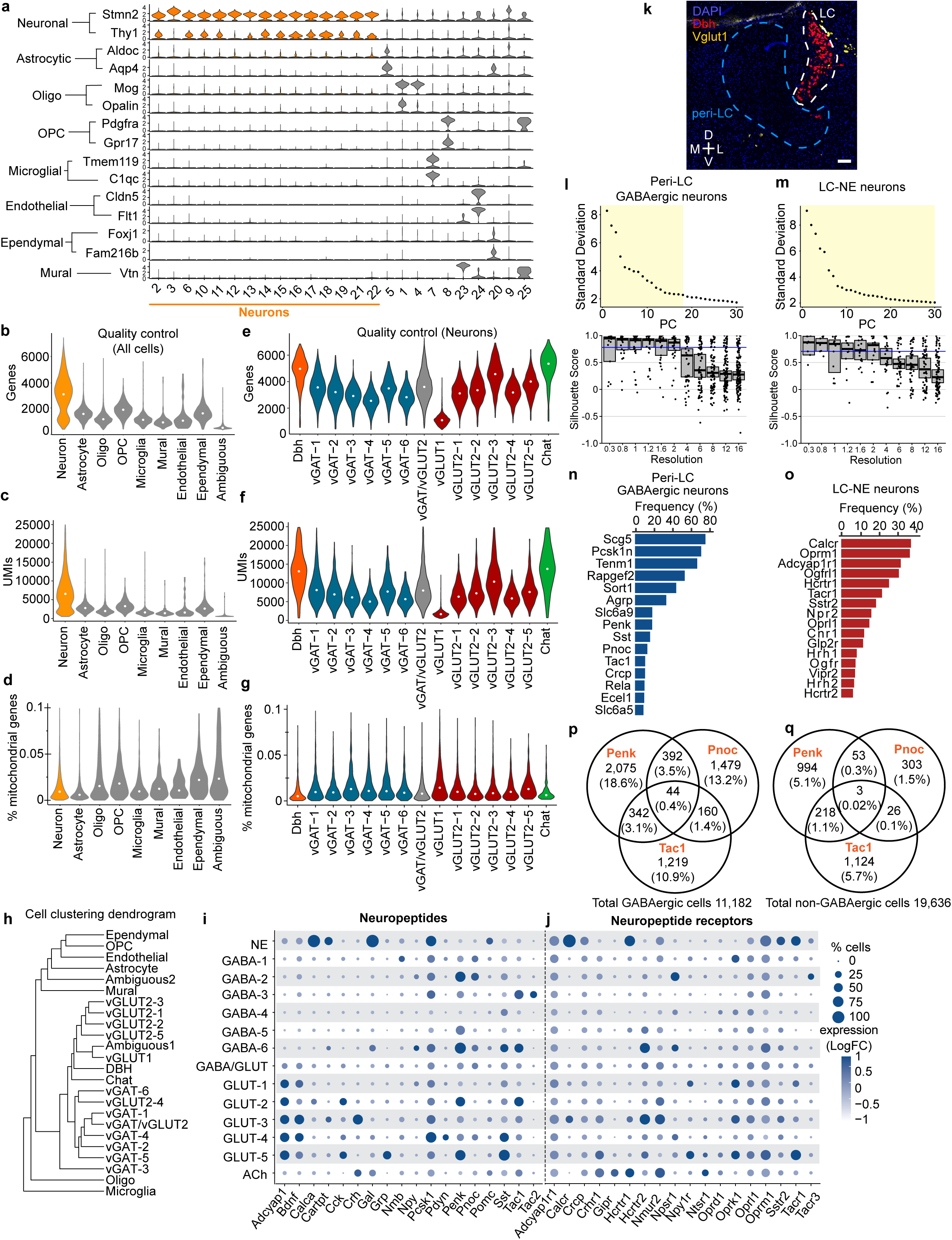
Single-cell RNA-sequencing of locus coeruleus and pericoerulear region. **(a)** Violin plots show expression of canonical marker genes in cells clustered by PCA based on transcriptional profile. The 15 neuronal clusters are shown in orange. **(b)** Genes detected per cell type. **(c)** Unique molecular identifiers (UMIs) per cell type. **(d)** Percent mitochondrial genes per cell type. **(e)** Genes detected per neuronal cell type. **(f)** Unique molecular identifiers (UMIs) per neuronal cell type. **(g)** Percent mitochondrial genes per neuronal cell type. **(h)** Cluster tree shows relationships between clustered cell types. **(i,j)** Dot plots of n=12,278 neurons are shown, with neuronal clusters on the y-axis and gene transcripts on the x-axis. Size of circles corresponds to percent of cells in the cluster expressing a specific transcript, while color intensity corresponds to the relative expression level of the transcript. Plots correspond to transcripts related to neuropeptides (**i**) and neuropeptide receptors (**j**). **(k)** Representative image for *in situ* hybridization of *Dbh* and vGLUT1 (*Slc17a7*) mRNA. Scale bar is 100 μm. **(l,m)** (Top) elbow plots show the number of principal components (PC) versus standard deviation and (bottom) silhouette scores show co-clustering performance for (**l**) peri-LC inhibitory neurons (Slc32a1, Gad1, Gad2) and **(m)** LC^NE^ neurons (Dbh, Th). Dots represent clusters at each resolution; boxes represent median scores with 95% confidence interval. **(n)** Graph shows the frequency of neuropeptide-expressing peri-LC inhibitory neurons. **(o)** Graph shows the frequency of neuropeptide receptor-expressing LC^NE^ neurons. **(p-q)** Venn diagrams show the overlap of Penk-, Pnoc-, and Tac1-expressing cells among cells that express (p) or do not express (q) GABAergic markers.

**Extended Data Fig. 4.**
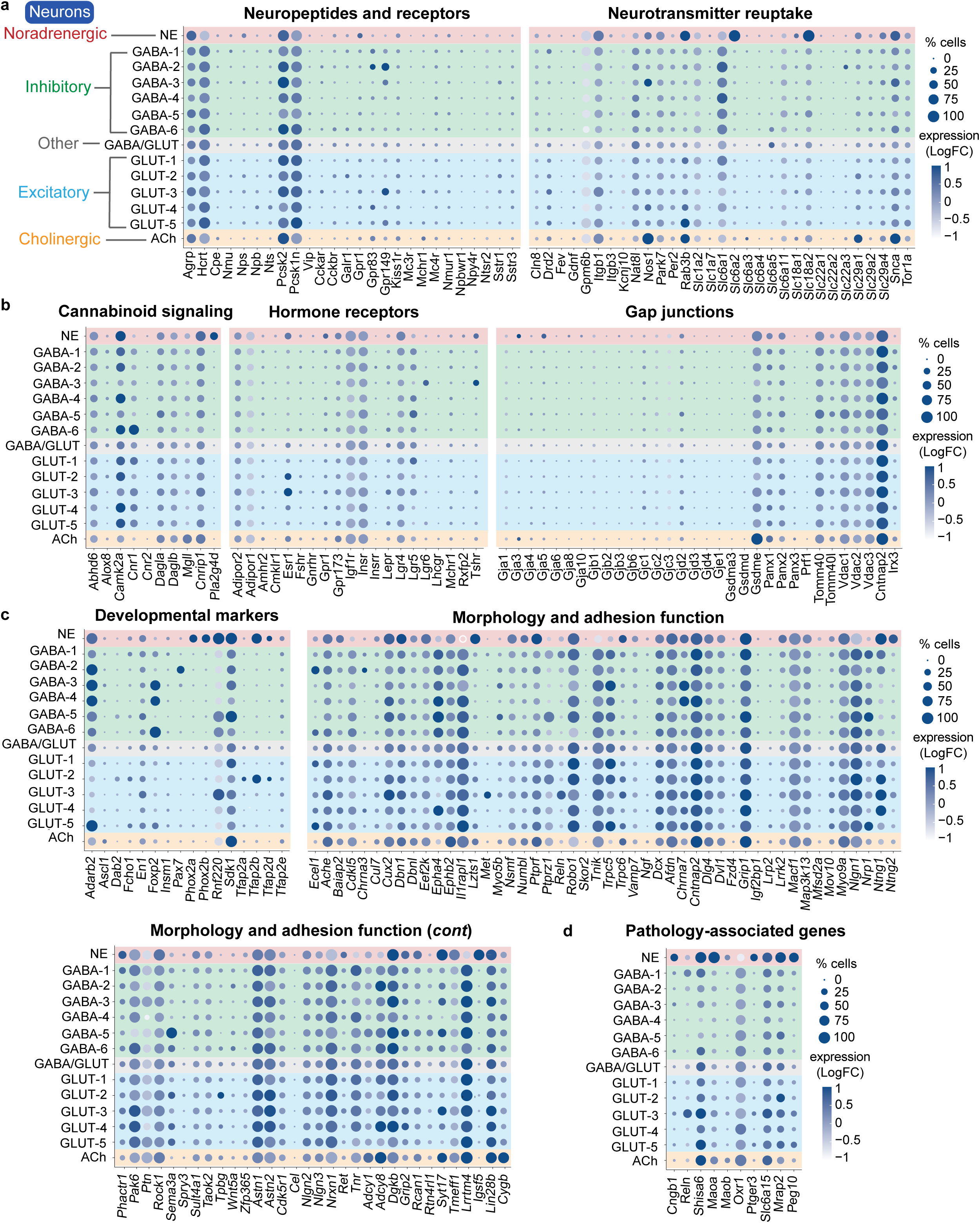
Neuromodulatory, signaling, structural, and disease-related genes in single-cell RNA sequencing of all peri-LC neurons (n=12,278). **(a)** Dot plot of neuropeptide and receptor (left), and neurotransmitter reuptake (right) genes across all neuron clusters. Circle size corresponds to percent of cells in the cluster expressing the specific transcript, while color intensity corresponds to its relative expression. **(b)** Dot plot of genes associated with cannabinoid signaling (left), hormone signaling (middle), and gap junctions (right) across all neuron clusters. **(c)** Dot plot of genes associated with development (left), and morphology and adhesion functions (right) across all neuron clusters. **(d)** Dot plot of genes associated with pathology across all neuron clusters.

**Extended Data Fig. 5.**
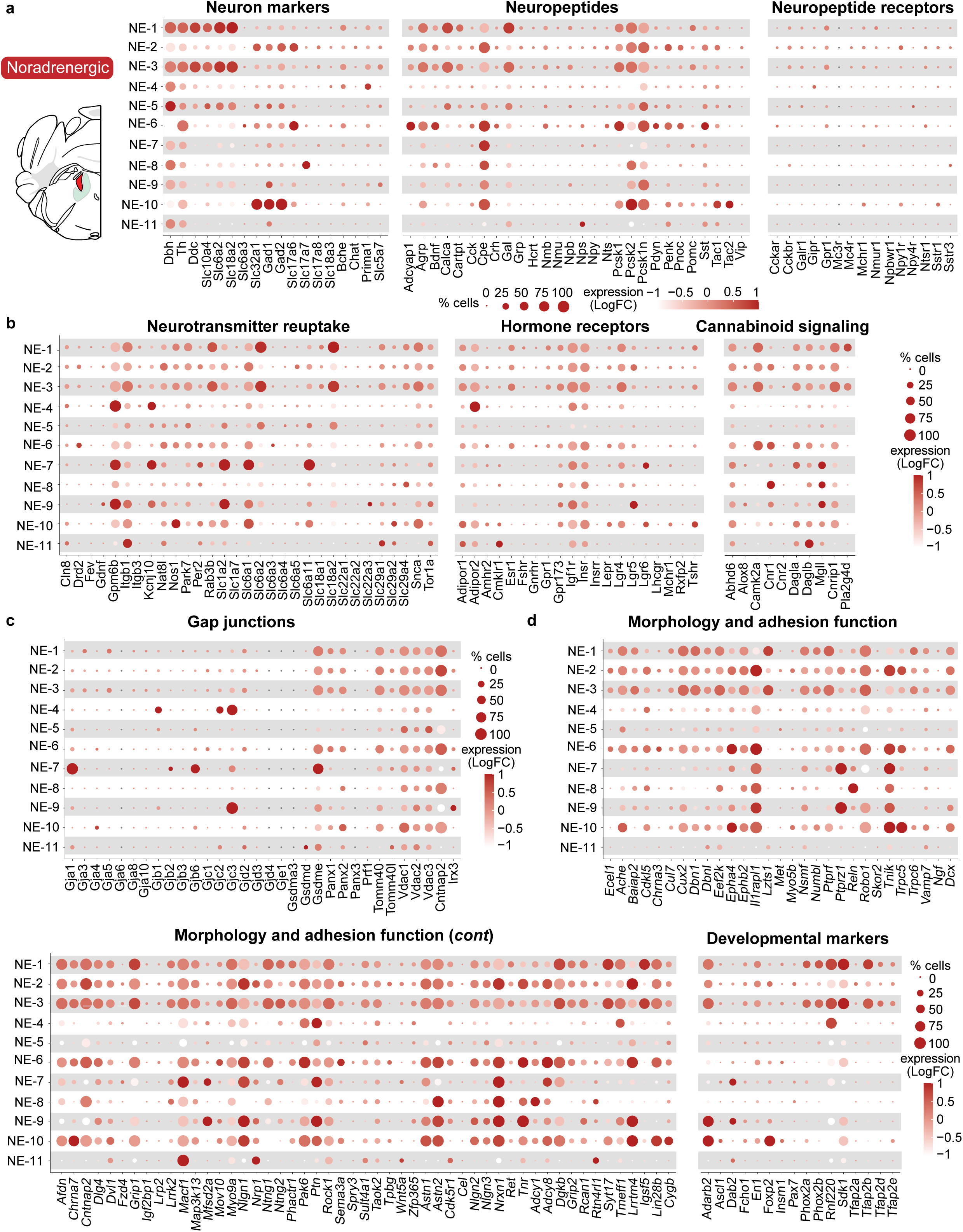
Neuromodulatory, signaling, and structural genes in single-cell RNA sequencing of LC^NE^ neurons (n=1,324). **(a)** Cartoon of the sampled locus coeruleus noradrenergic neurons (left). Dot plot of neuronal subtype (left), neuropeptide (middle), and receptor (right) genes across LC^NE^ neuron clusters. Circle size corresponds to percent of cells in the cluster expressing the specific transcript, while color intensity corresponds to its relative expression. **(b)** Dot plot of genes associated with neurotransmitter reuptake (left), hormone signaling (middle), and cannabinoid signaling (right) across LC^NE^ neuron clusters. **(c)** Dot plot of genes associated with gap junctions across LC^NE^ neuron clusters. **(d)** Dot plot of genes associated with morphology and adhesion functions and development across LC^NE^ neuron clusters.

**Extended Data Fig. 6.**
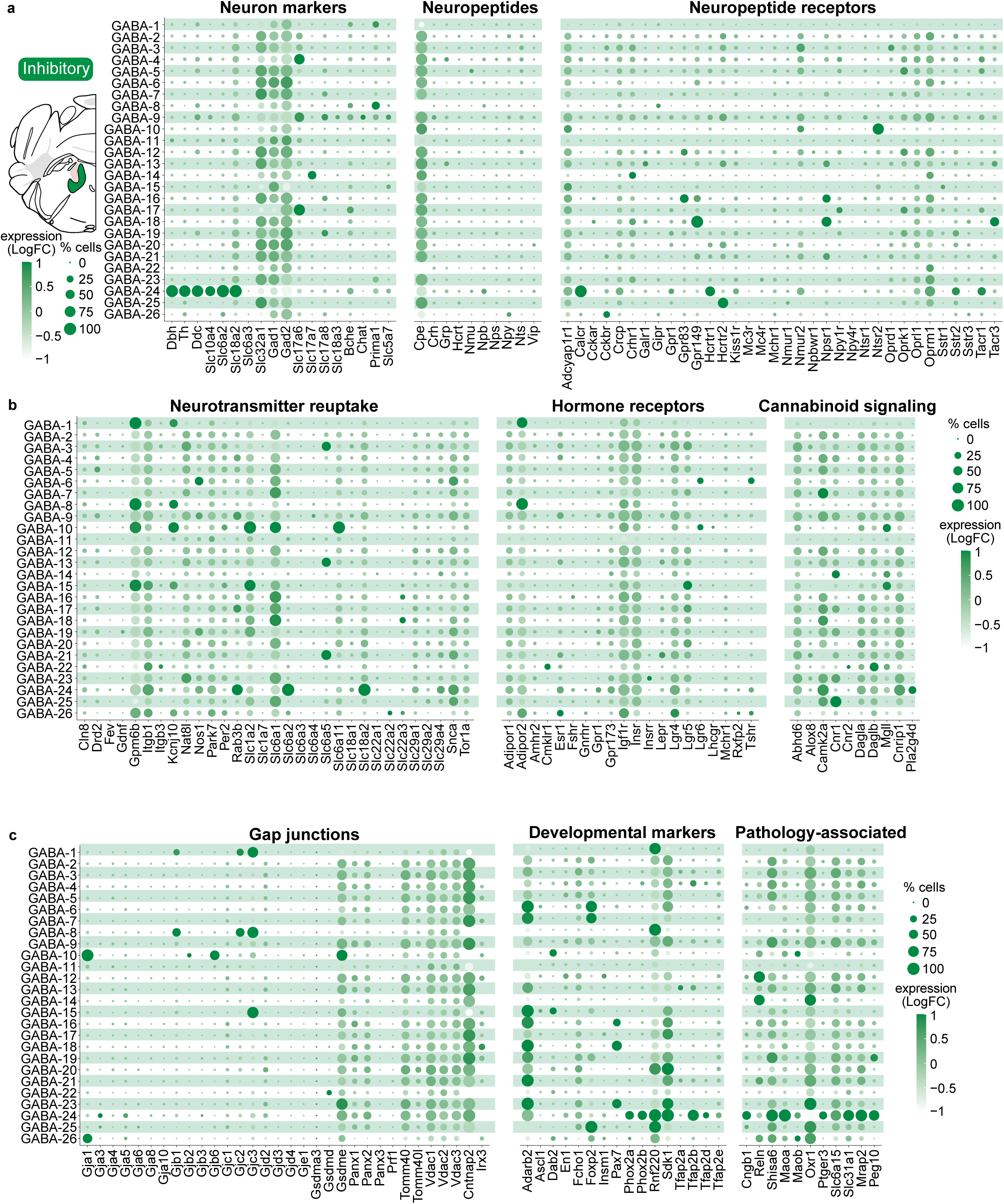
Neuromodulatory, signaling, and structural genes in single-cell RNA sequencing of peri-LC^GABA^ neurons (n=11,182). **(a)** Cartoon of the sampled peri-LC inhibitory neurons (left). Dot plot of neuronal subtype (left), neuropeptide (middle), and receptor (right) genes across peri-LC^GABA^ neuron clusters. Circle size corresponds to percent of cells in the cluster expressing the specific transcript, while color intensity corresponds to its relative expression. **(b)** Dot plot of genes associated with neurotransmitter reuptake (left), hormone signaling (middle), and cannabinoid signaling (right) across peri-LC^GABA^ neuron clusters. **(c)** Dot plot of genes associated with gap junctions across peri-LC^GABA^ neuron clusters.

**Extended Data Fig. 7.**
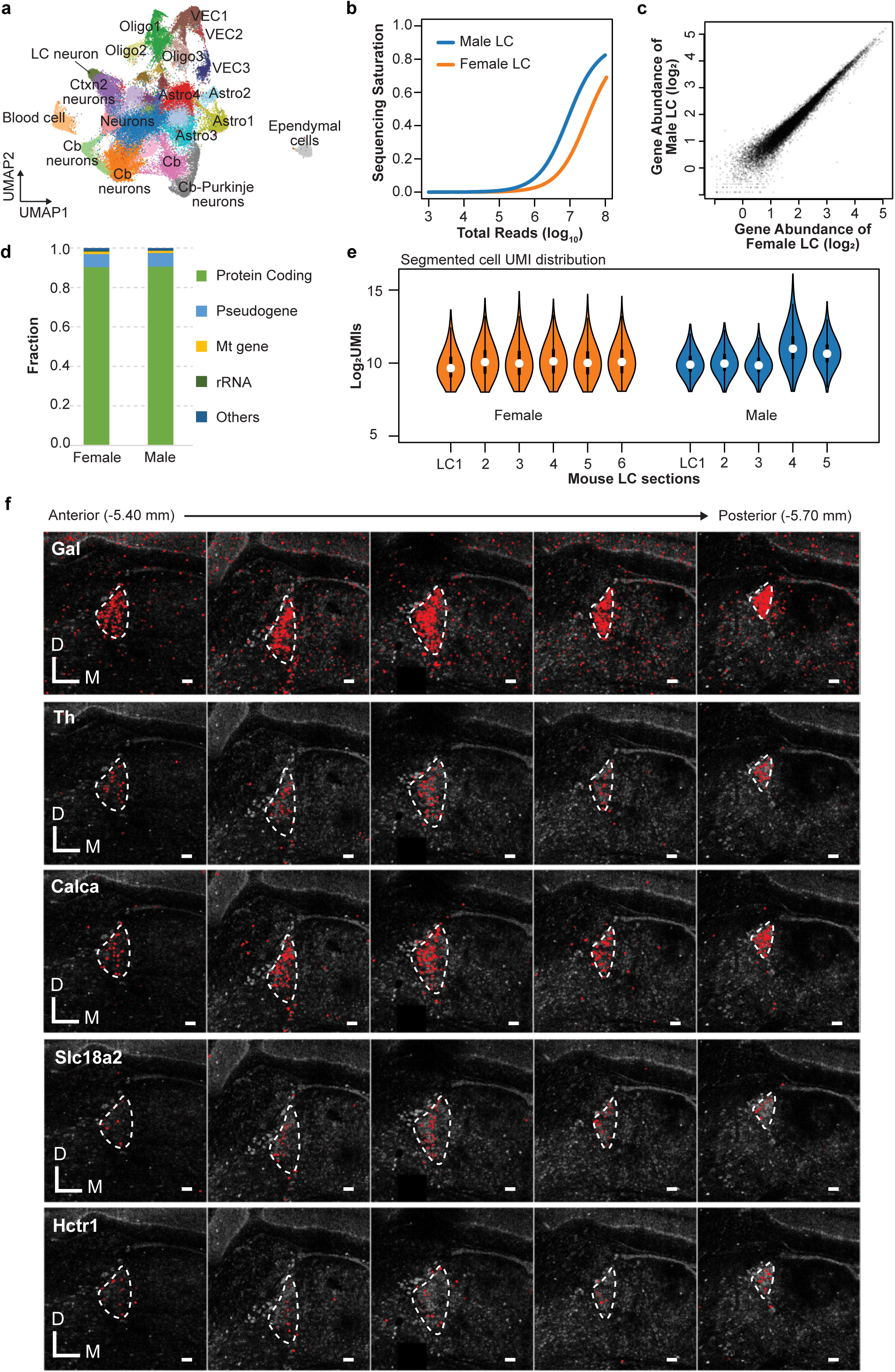
PIXEL-seq quality control. **(a)** Uniform Manifold Approximation and Projection (UMAP) embedding of ∼59,000 cells passing quality control metrics, colored by clustering analysis. **(b)** Sequencing reads and saturation for 11 brain slices (6 female, 5 male) containing the LC use for PIXEL-seq. **(c)** Comparison of transcripts found in brain slices of male and female mice used for PIXEL-seq. **(d)** Proportion of the RNA type in the brain slices obtained from male and female mice used for PIXEL-seq. **(e)** Unique molecular identifier (UMI) counts for the 11 brain slices used for PIXEL-seq. **(f)** Spatial expressions of *Gal, Th, Calca, Slc18a2, and Hctr1* in the LC region are shown along the anterior-posterior axis (Bregma -5.40 mm to -5.70 mm). Scale bars are 100 μm.

**Extended Data Fig. 8.**
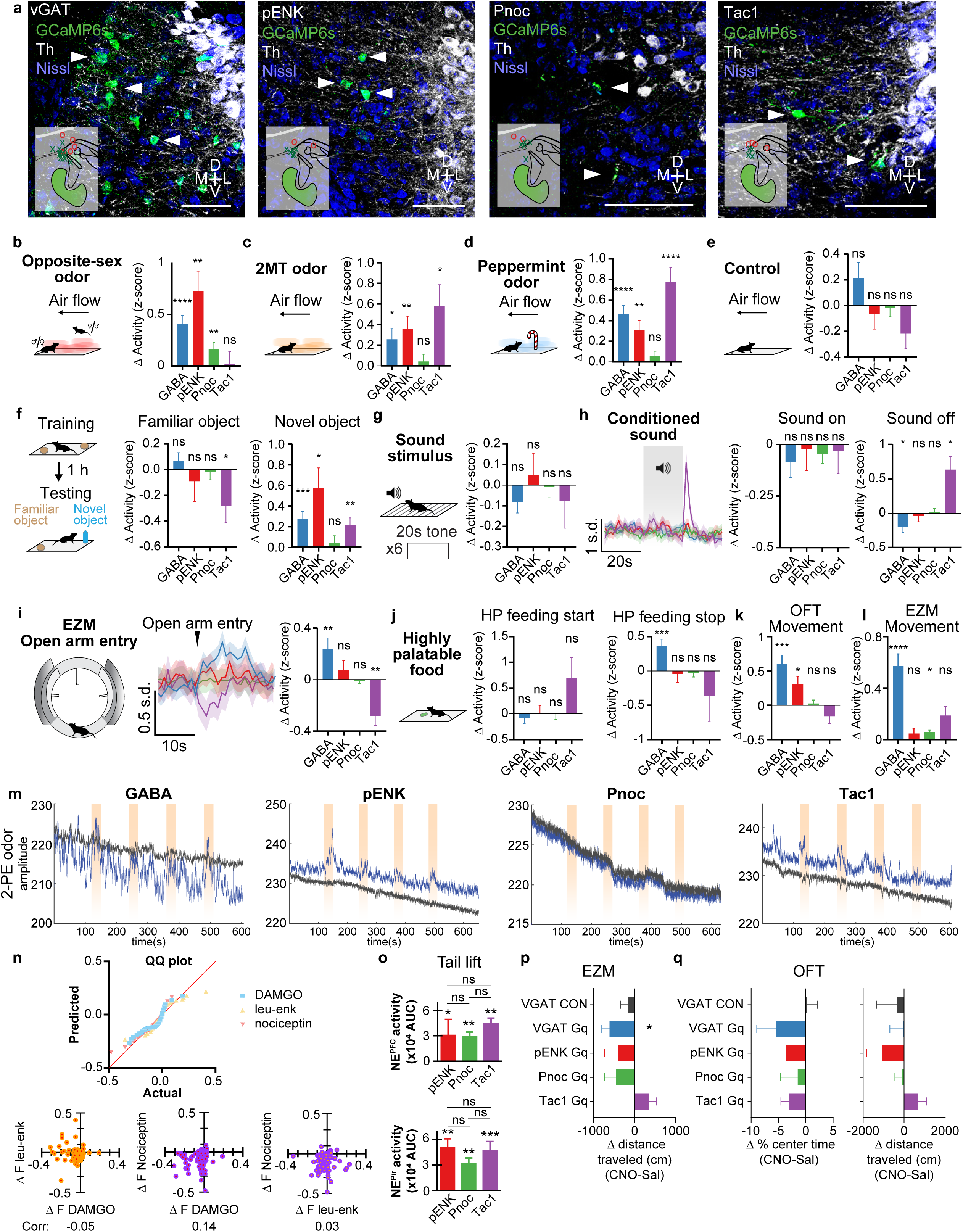
Peri-LC neuronal subpopulations differentially respond to arousing and stress stimuli. **(a)** Example images of GCaMP6s expression in peri-LC subpopulations. Insets show placements of fibers, with ‘x’ representing on-target fibers and expression and ‘o’ representing no or off-target expression. Scale bars is 50 μm. **(b)** Peri-LC neurons are modulated by 30 s exposure to odor of opposite-sex mice. **(c)** Peri-LC neurons are modulated by 30 s exposure to 2MT. **(d)** Peri-LC neurons are modulated by 30 s exposure to peppermint oil odor. **(e)** Peri-LC neurons are not affected by 30 s of blowing air through chamber. **(f)** Peri-LC neurons are modulated by interactions with a novel object or a familiar object. **(g)** Peri-LC neurons are unaffected by exposure to a tone that had not yet been paired with a shock. **(h)** Peri-LC neurons do not respond to a conditioned tone, but GABA neurons decrease activity and Tac1 neurons increase activity after the tone stops. **(i)** Peri-LC GABA neurons increase activity and Tac1 neurons decrease activity when an animal enters the open arm of an elevated zero maze. **(j)** Peri-LC^GABA^ neurons increase activity after they stop eating highly palatable food; no other conditions are affected. Bars represent change in activity before vs. after events. Periods of comparison were 30s for odor experiments and air flow, 20s for sound stimuli, 10s for feeding, and 5s for novel/familiar object interaction and maze exploration. **(k)** Activity of peri-LC neurons during movement in the open field test. Bars represent difference in activity during movement vs. non-movement for each trial. **(l)** Activity of peri-LC neurons during movement in the elevated-zero maze. Bars represent difference in activity during movement vs. non-movement for each trial. **(m)** Representative signal in 470nm (blue) and 405nm (gray) channels for peri-LC GABA, pENK, Pnoc, and Tac1 recordings of 2-PE odor. *p< 0.05, ** p<0.01, ***p<0.001, ****p<0.0001 by paired two-tailed *t*-test. Error bars indicate SEM. n=12 animals for GABA photometry, 6 for pENK, 5 for Pnoc, and 4 for Tac1. **(n)** Top, QQ plot shows divergence from test normality of LC-NE neuronal responses for the three peptide perfusions. Bottom, correlational plots of LC-NE responses for combinations of peptides, with the Pearson’s correlational coefficient shown on the bottom. **(o)** Area under curve over 10s of photometry recordings of NE activity (as measured by GRAB_NE2m) in the PFC (top) and Pir (bottom) during tail lifts in the pENK-Cre (N=6), Pnoc-Cre (N=5), and Tac1-Cre (N=8) mice used for the peri-LC photostimulation experiments. *p<0.05, **p<0.01, ***p<0.001. **(p)** The difference in total distance traveled in the elevated zero maze calculated as the full 20 min of assay ∼30 min after saline or 1 mg/kg CNO treatment are shown for VGAT fluorophore control (N=7), VGAT Gq (N=7), pENK Gq (N=10), Pnoc Gq (N=9), and Tac1 Gq (N=13) mice. *p<0.05. **(q)** The difference in time spent in the center and total distance traveled (20 min of assay 30 min after saline or 1 mg/kg CNO treatment) for mice in open field test are shown for VGAT mCherry control (N=7), VGAT Gq (N=7), Penk Gq (N=10), Pnoc Gq (N=9), and Tac1 Gq (N=13) mice.

**Extended Data Table 1.** Electrophysiological properties of recorded LC and peri-LC GABAergic neurons.

|  |  |  |  |
| --- | --- | --- | --- |
| **LC<sup>NE</sup> neurons** |  |  |  |
| | IPSC amplitude | $53.8 \pm 7.3$ pA | n/N = 18/9 |
| | Paired pulse ratio (50 ms IPI) | $1.13 \pm 0.11$ | n/N = 16/9 |
| | Rise time | $3.15 \pm 0.47$ ms | n/N = 18/9 |
| | Width | $16.1 \pm 1.7$ ms | n/N = 18/9 |
| | Decay constant | $17.2 \pm 1.2$ ms | n/N = 18/9 |
| **Peri-LC GABA neurons** |  |  |  |
| | Capacitance | $27.5 \pm 1.1$ pF | n/N = 24/12 |
| | Membrane resistance | $316.3 \pm 29.8$ mΩ | n/N = 23/12 |
| | Resting membrane potential | $-53.5 \pm 1.4$ mV | n/N = 13/7 |
| | Basal firing rate | $3.5 \pm 1.0$ Hz | n/N = 6/5 |
| | Firing rate +200 pA | $11.6 \pm 1.5$ Hz | n/N = 11/7 |
| | Firing rate +400 pA | $23.2 \pm 2.3$ Hz | n/N = 11/7 |

## Notes

### Summary of Updates

Figures 5 and S8 revised to include new behavioral and fiber photometry data; results clarified with new language

