## Supplementary Materials - Pathology, Neurotransmission, and GPCR-related genes for "A diverse population of pericoerulear neurons controls arousal and exploratory behaviors"

### A diverse network of pericoerulear neurons control arousal states

**This PDF file includes:**

Figures S1-S6

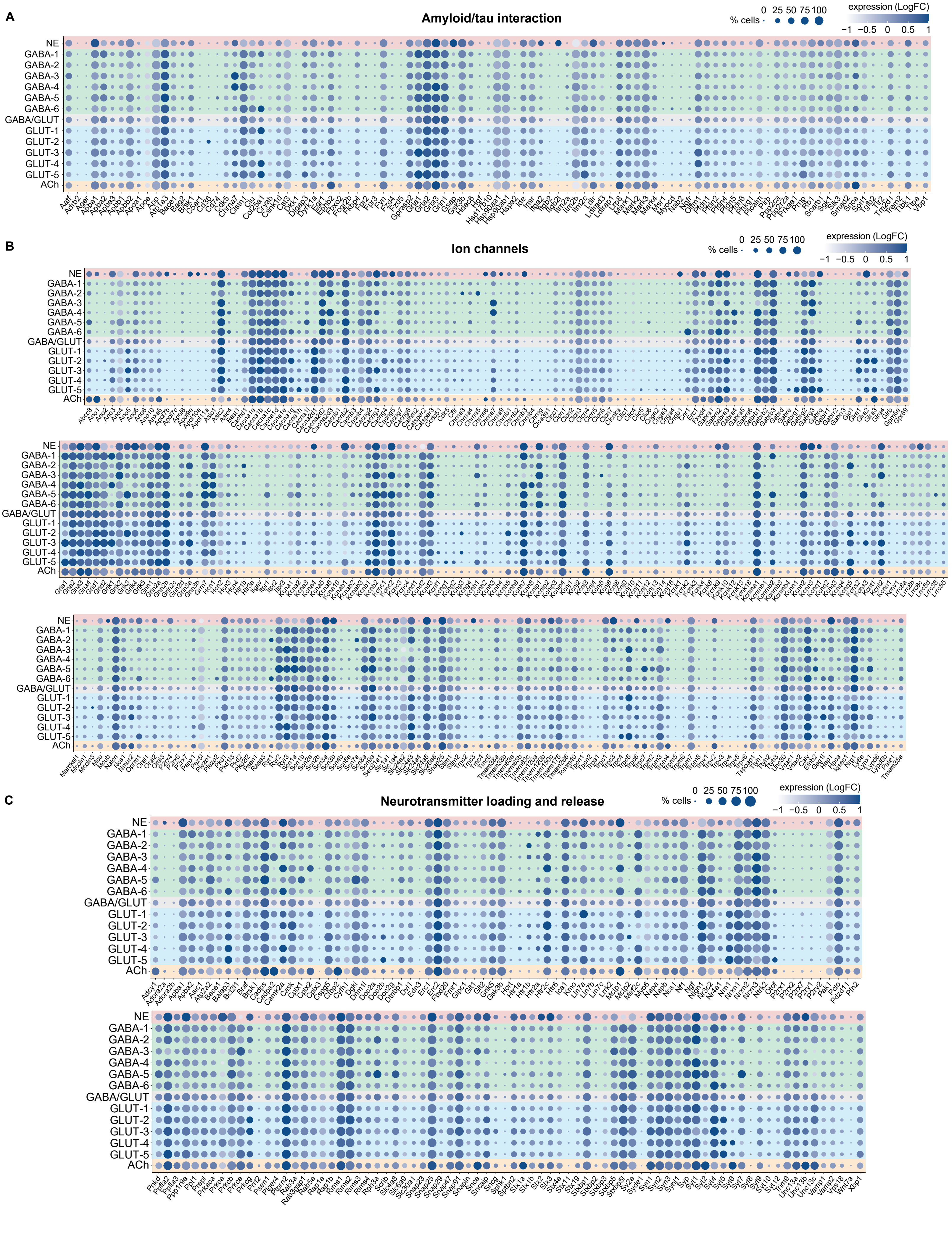
A
Amyloid/tau interaction
B
Ion channels
C
Neurotransmitter loading and release
% cells
0 25 50 75 100
expression (LogFC)
−1 −0.5 0 0.5 1
NE
GABA-1
GABA-2
GABA-3
GABA-4
GABA-5
GABA-6
GABA/GLUT
GLUT-1
GLUT-2
GLUT-3
GLUT-4
GLUT-5
ACh

**Supplementary Fig. 1. Additional disease-related, ion channel, and neurotransmitter genes in single-cell RNA sequencing of all peri-LC neurons.**

**(A)** Dot plot of genes associated with amyloid and tau interaction across all neuron clusters. Circle size corresponds to percent of cells in the cluster expressing the specific transcript, while color intensity corresponds to its relative expression.

**(B)** Dot plot of genes associated with ion channels across all neuron clusters.

**(C)** Dot plot of genes associated with neurotransmitter loading and release across all neuron clusters.

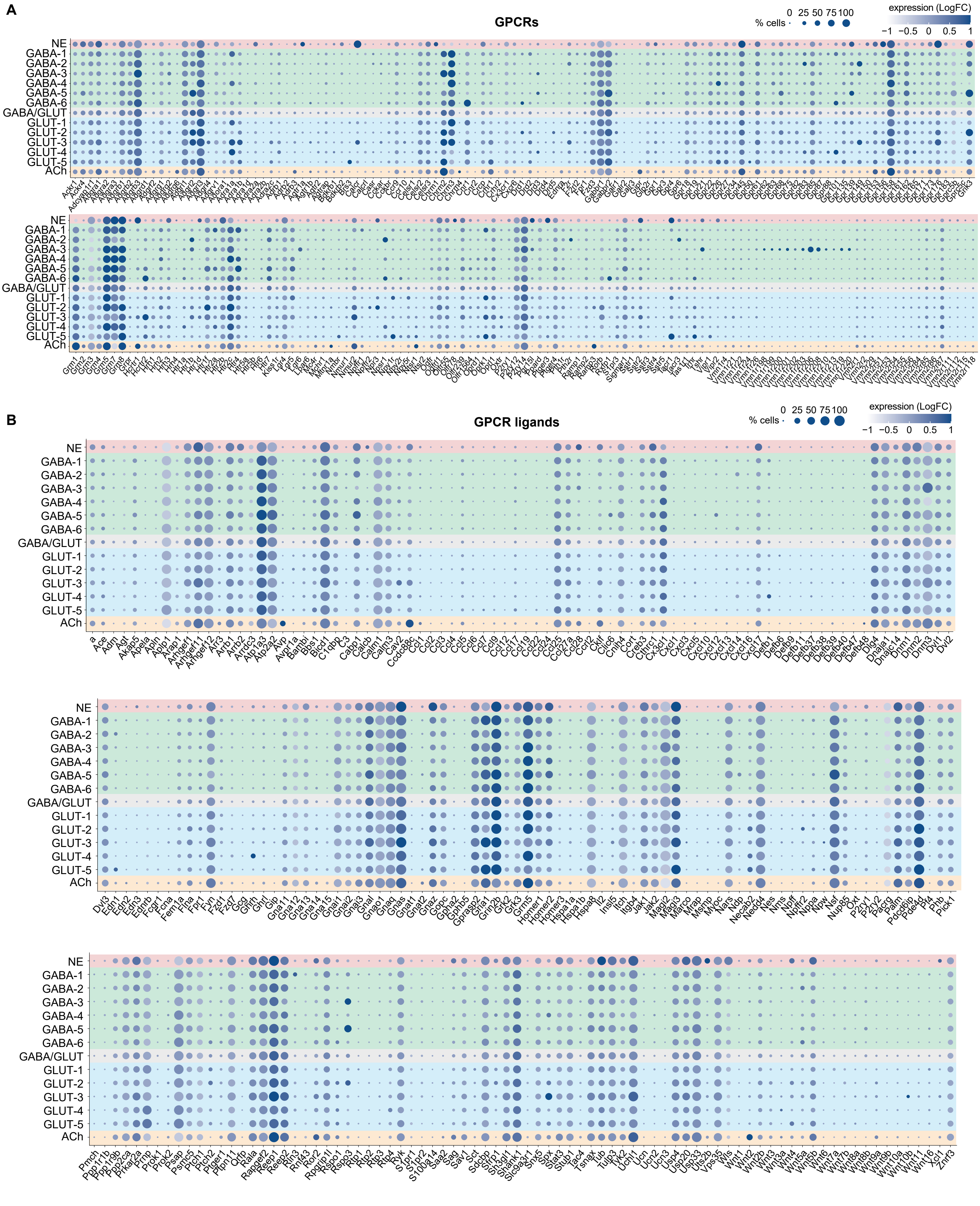
A
GPCRs
% cells
0 25 50 75 100
expression (LogFC)
−1 −0.5 0 0.5 1
NE
GABA-1
GABA-2
GABA-3
GABA-4
GABA-5
GABA-6
GABA/GLUT
GLUT-1
GLUT-2
GLUT-3
GLUT-4
GLUT-5
ACh
B
GPCR ligands

**Supplementary Fig. 2. G-protein coupled receptor (GPCR) and ligand genes in single-cell RNA sequencing of all peri-LC neurons.**

**(A)** Dot plot of genes associated with GPCRs across all neuron clusters. Circle size corresponds to percent of cells in the cluster expressing the specific transcript, while color intensity corresponds to its relative expression.

**(B)** Dot plot of genes associated with GPCR ligands across all neuron clusters.

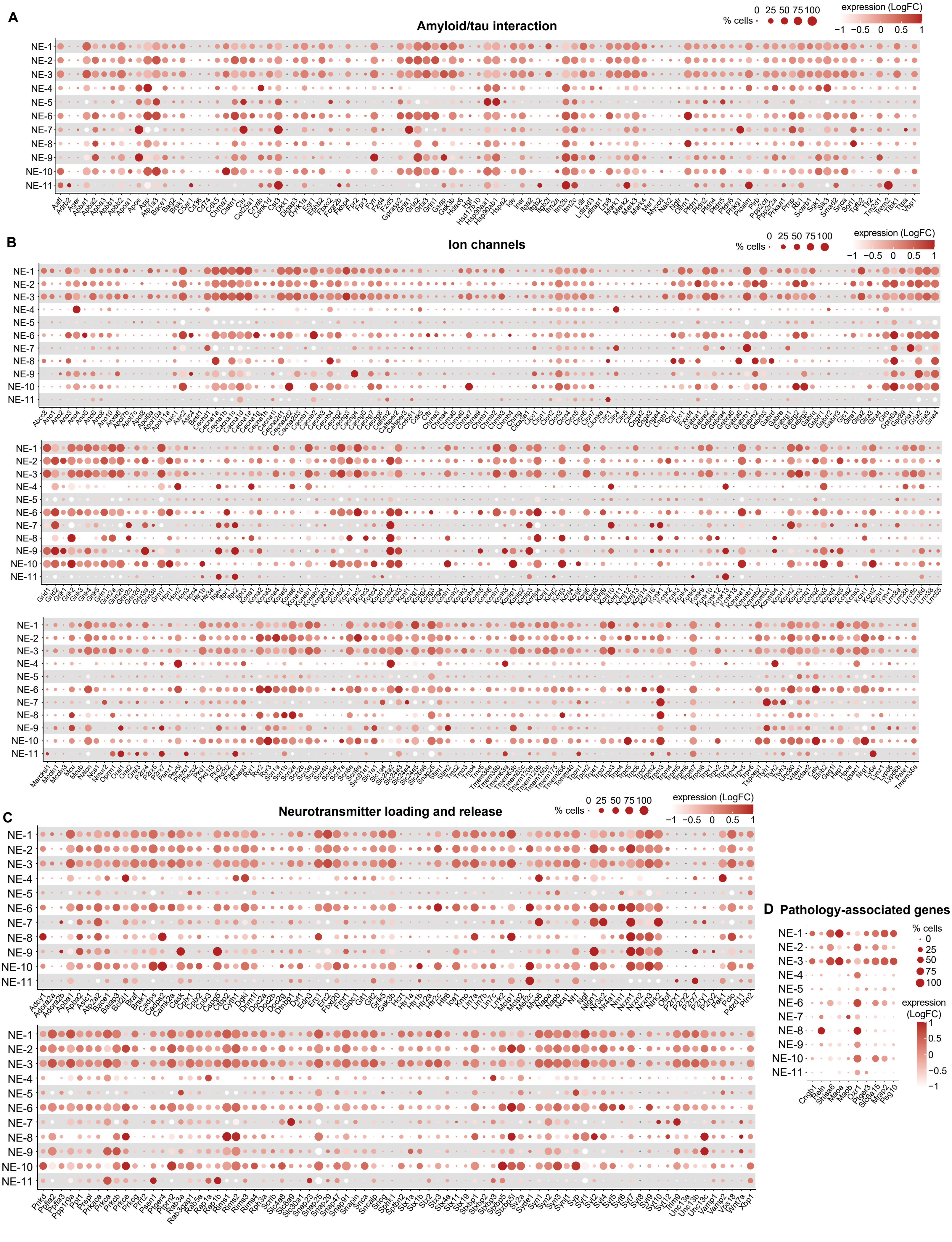
A
Amyloid/tau interaction
% cells
0 25 50 75 100
expression (LogFC)
−1 −0.5 0 0.5 1
B
Ion channels
C
Neurotransmitter loading and release
D
Pathology-associated genes
NE-1
NE-2
NE-3
NE-4
NE-5
NE-6
NE-7
NE-8
NE-9
NE-10
NE-11

**Supplementary Fig. 3. Disease-related, ion channel, and neurotransmitter genes in single-cell RNA sequencing of $LC^{NE}$ neurons.**

**(A)** Dot plot of genes associated with amyloid and tau interaction across $LC^{NE}$ neuron clusters. Circle size corresponds to percent of cells in the cluster expressing the specific transcript, while color intensity corresponds to its relative expression.

**(B)** Dot plot of genes associated with ion channels across $LC^{NE}$ neuron clusters.

**(C)** Dot plot of genes associated with neurotransmitter loading and release across $LC^{NE}$ neuron clusters.

**(D)** Dot plot of genes associated with pathology across $LC^{NE}$ neuron clusters.

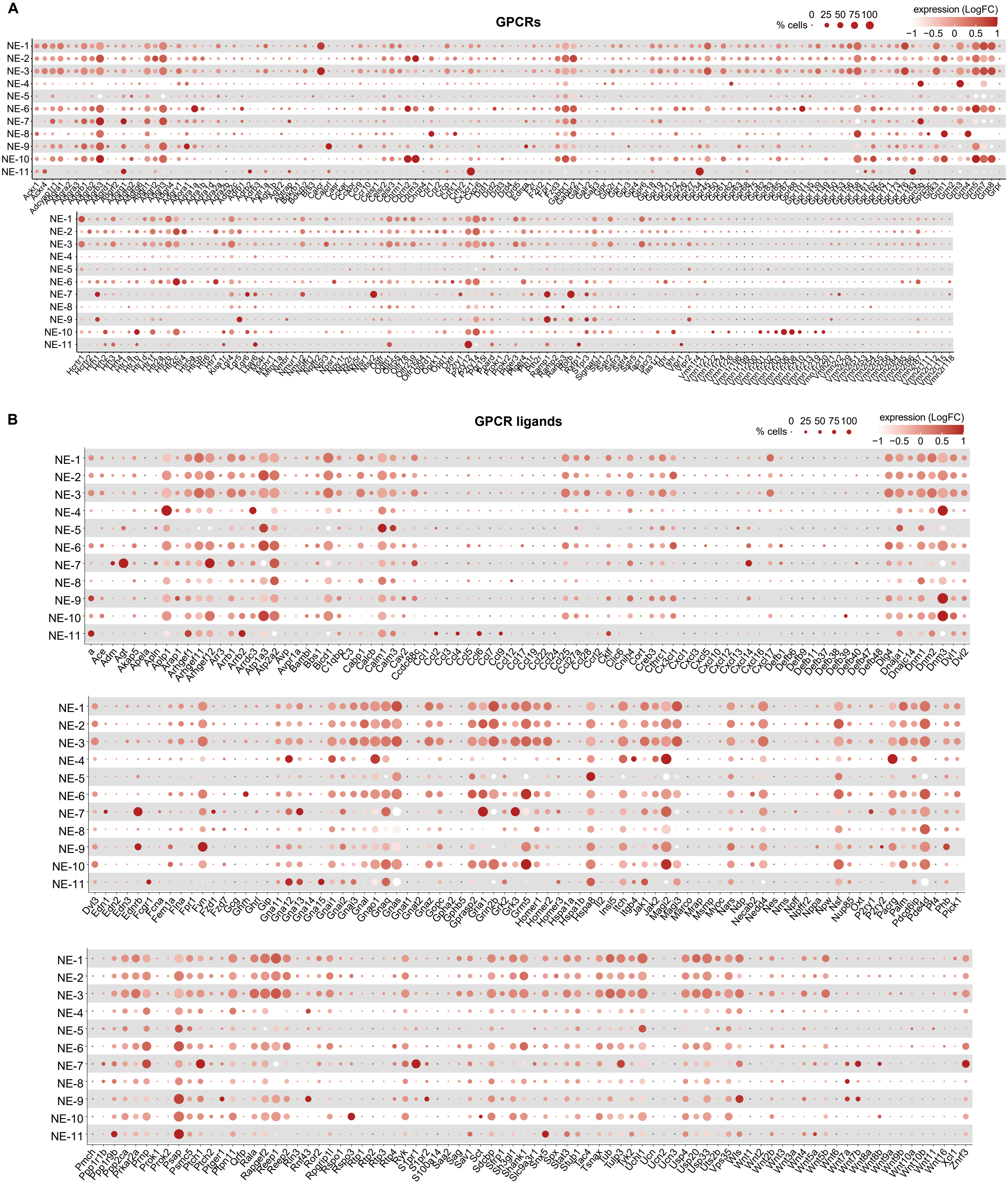
A
GPCRs
% cells
0 25 50 75 100
expression (LogFC)
−1 −0.5 0 0.5 1
NE-1
NE-2
NE-3
NE-4
NE-5
NE-6
NE-7
NE-8
NE-9
NE-10
NE-11
B
GPCR ligands
% cells
0 25 50 75 100
expression (LogFC)
−1 −0.5 0 0.5 1

**Supplementary Fig. 4. G-protein coupled receptor (GPCR) and ligand genes in single-cell RNA sequencing of $LC^{NE}$ neurons.**
**(A)** Dot plot of genes associated with GPCRs across $LC^{NE}$ neuron clusters. Circle size corresponds to percent of cells in the cluster expressing the specific transcript, while color intensity corresponds to its relative expression.
**(B)** Dot plot of genes associated with GPCR ligands across $LC^{NE}$ neuron clusters**.**

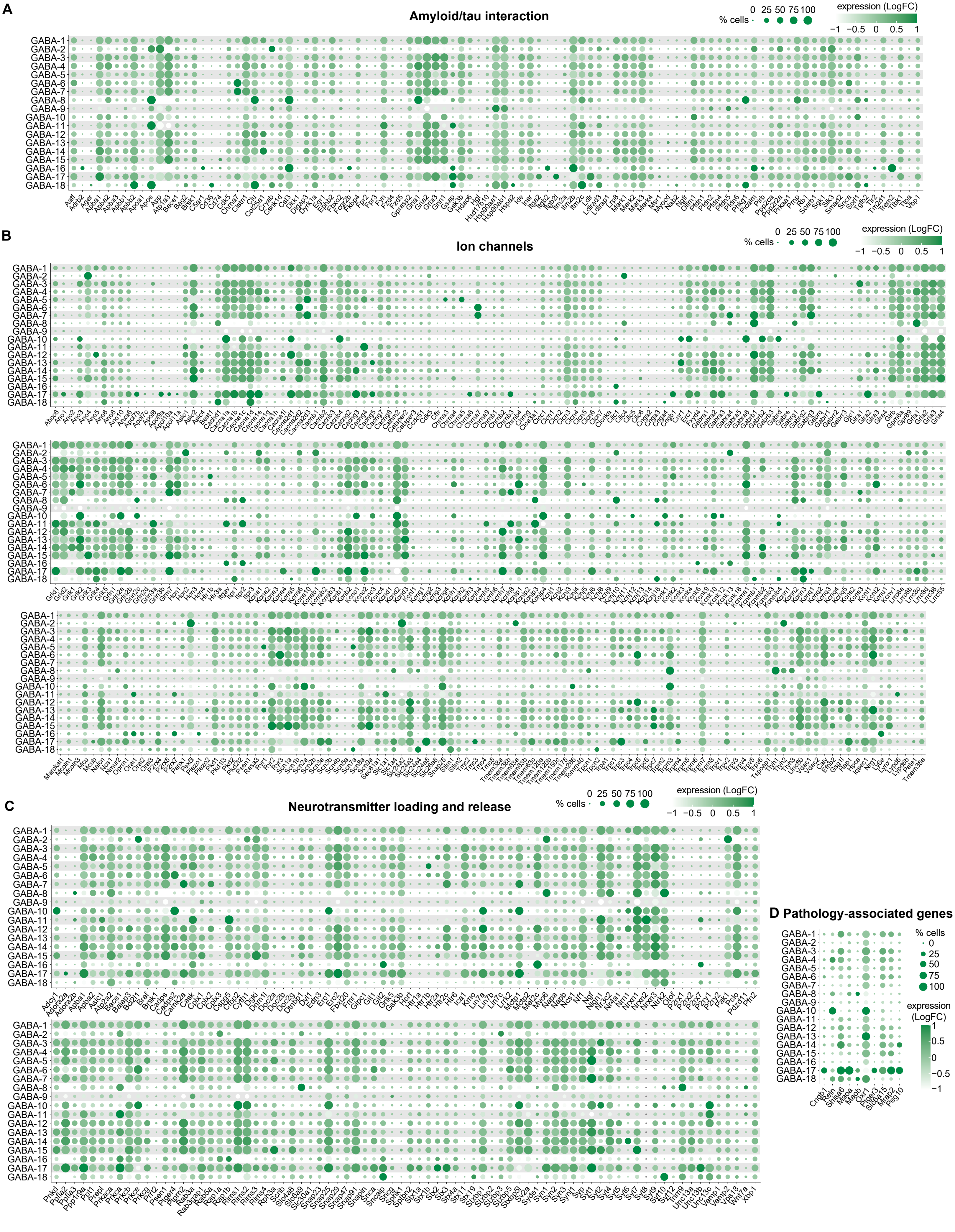
A
Amyloid/tau interaction
% cells
0 25 50 75 100
expression (LogFC)
−1 −0.5 0 0.5 1
B
Ion channels
C
Neurotransmitter loading and release
D Pathology-associated genes
GABA-1
GABA-2
GABA-3
GABA-4
GABA-5
GABA-6
GABA-7
GABA-8
GABA-9
GABA-10
GABA-11
GABA-12
GABA-13
GABA-14
GABA-15
GABA-16
GABA-17
GABA-18
Cngb1
Reln
Snisa6
Maoa
Maob
Oxr1
Ptger3
Slc6a15
Mrap2
Peg10

**Supplementary Fig. 5. Disease-related, ion channel, and neurotransmitter genes in single-cell RNA sequencing of peri-LC$^{GABA}$ neurons.**

**(A)** Dot plot of genes associated with amyloid and tau interaction across peri-LC$^{GABA}$ neuron clusters. Circle size corresponds to percent of cells in the cluster expressing the specific transcript, while color intensity corresponds to its relative expression.

**(B)** Dot plot of genes associated with ion channels across peri-LC$^{GABA}$ neuron clusters.

**(C)** Dot plot of genes associated with neurotransmitter loading and release across peri-LC$^{GABA}$ neuron clusters.

**(D)** Dot plot of genes associated with pathology across peri-LC$^{GABA}$ neuron clusters.

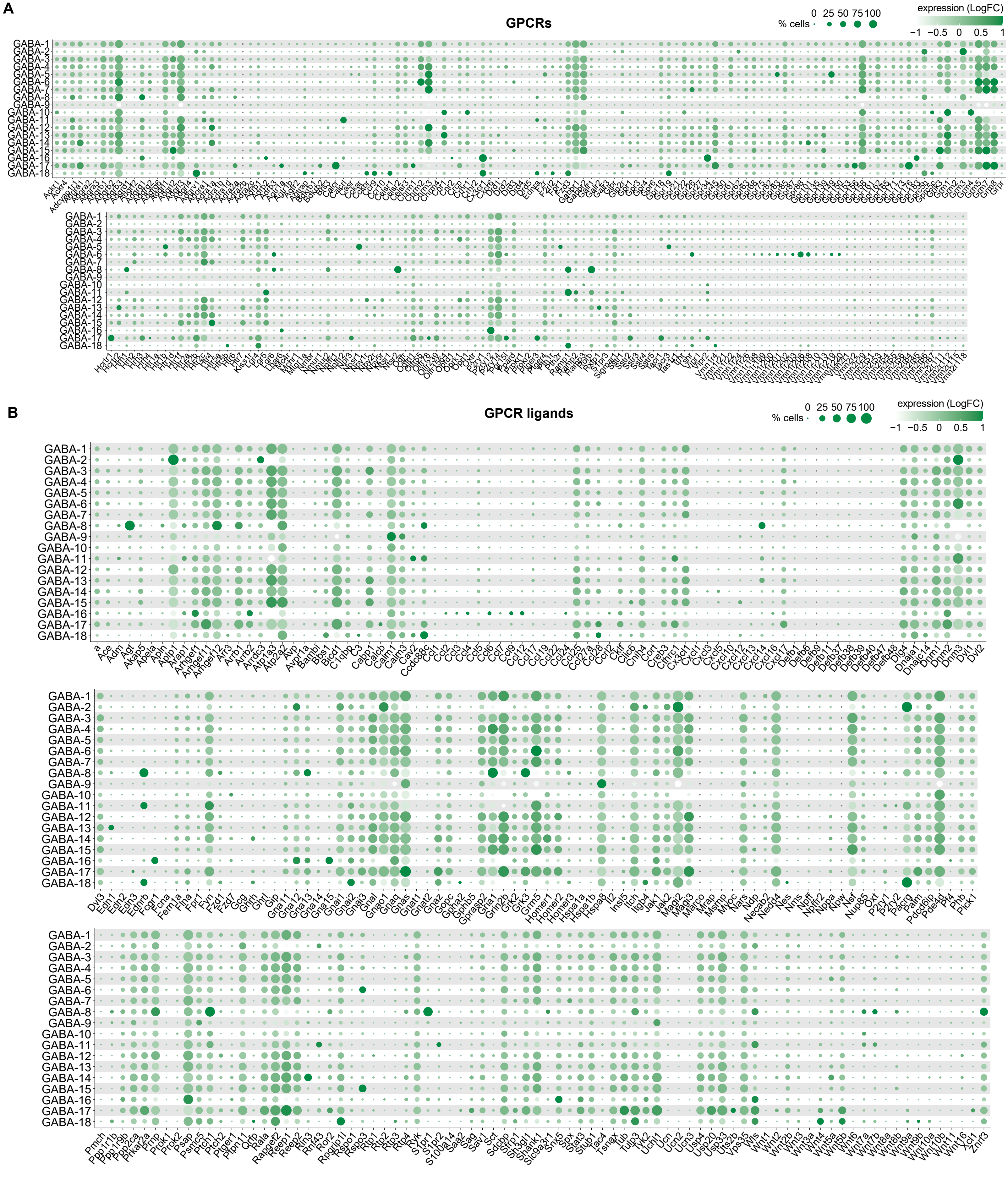
A
GPCRs
% cells
0 25 50 75 100
expression (LogFC)
−1 −0.5 0 0.5 1
GABA-1
GABA-2
GABA-3
GABA-4
GABA-5
GABA-6
GABA-7
GABA-8
GABA-9
GABA-10
GABA-11
GABA-12
GABA-13
GABA-14
GABA-15
GABA-16
GABA-17
GABA-18
B
GPCR ligands
% cells
0 25 50 75 100
expression (LogFC)
−1 −0.5 0 0.5 1

**Supplementary Fig. 6. G-protein coupled receptor (GPCR) and ligand genes in single-cell RNA sequencing of peri-LC$^{GABA}$ neurons.**

**(A)** Dot plot of genes associated with GPCRs across peri-LC$^{GABA}$ neuron clusters. Circle size corresponds to percent of cells in the cluster expressing the specific transcript, while color intensity corresponds to its relative expression.

**(B)** Dot plot of genes associated with GPCR ligands across peri-LC$^{GABA}$ neuron clusters**.**
